# Aberrant Molecular Myelin Architecture in Charcot-Marie-Tooth Disease Type 1A and Hereditary Neuropathy with Liability to Pressure Palsies

**DOI:** 10.1101/2024.05.10.592618

**Authors:** Kathryn R. Moss, Marvis A. Arowolo, Dave R. Gutierrez, Ahmet Höke

**Author notes:** Correspondence: Kathryn R. Moss.

## Abstract

Charcot-Marie-Tooth Disease Type 1A (CMT1A) and Hereditary Neuropathy with Liability to Pressure Palsies (HNPP) are the most common inherited peripheral neuropathies and arise from copy number variation of the *Peripheral Myelin Protein 22* gene (*PMP22*). While secondary axon degeneration has been proposed as a primary driver of pathogenesis, our prior work demonstrated neuromuscular deficits in CMT1A mice in the absence of overt axonal loss, prompting investigation into primary myelin dysfunction. Here, we reveal that altered *PMP22* dosage profoundly disrupts molecular architecture at critical myelin subdomains, Schmidt-Lanterman incisures (SLIs) and Nodes of Ranvier. Using high-resolution confocal imaging of teased peripheral nerve fibers from CMT1A and HNPP model mice, we identified disorganization of adherens junctions, mislocalization of Connexin29, and altered distribution of nodal ion channels in CMT1A and HNPP, with several defects more pronounced in CMT1A, aligning with clinical severity. Notably, Kv1.2 and Caspr mislocalization along the internode and Nav nodal widening suggest disruption of axoglial domains essential for saltatory conduction. Together, these phenotypes support a model in which PMP22 governs myelin architecture, likely through adherens junction regulation, with its dysregulation predicted to impair metabolic support and axonal ion homeostasis, thereby compromising the structural and functional integrity of myelin and contributing directly to disease pathogenesis. These findings shift the pathogenic paradigm for CMT1A and HNPP from axonal degeneration to primary myelin failure and highlight actionable molecular targets for therapeutic intervention. This study offers mechanistic insight into CMT1A and HNPP and provides a conceptual framework with broad relevance to other dysmyelinating disorders.

**Main Points:**

- *PMP22* copy number variation disrupts myelin architecture at SLIs and Nodes of Ranvier.
- Adherens junction and axoglial domain defects are more severe in CMT1A than HNPP.
- Findings support primary myelin dysfunction as a key driver of pathogenesis.

## Introduction

Charcot-Marie-Tooth Disease Type 1A (CMT1A) and Hereditary Neuropathy with Liability to Pressure Palsies (HNPP) are the most common inherited peripheral neuropathies, together accounting for approximately 65% of genetically defined CMT cases^1^. The overall prevalence of CMT is estimated at 1 in 2,500 individuals, making it one of the most prevalent inherited neurological conditions^2^. Remarkably, CMT1A and HNPP result from opposing copy number variations of the same gene: *Peripheral Myelin Protein 22* (*PMP22*), which encodes PMP22 protein. Duplication of *PMP22* causes CMT1A, a progressive dysmyelinating neuropathy that typically presents in childhood and leads to steadily worsening motor and sensory deficits, often resulting in lifelong disability^3^. In contrast, deletion of *PMP22* causes HNPP, a milder disorder characterized by transient, focal neuropathies triggered by mechanical compression, with onset typically in adolescence or adulthood^4^.

Although recovery is common in HNPP, repeated episodes can cause permanent nerve damage^4^. Despite differences in severity and progression, both conditions are associated with significant morbidity, and no disease-modifying therapies currently exist^5,6^.

PMP22 is a Schwann cell-enriched membrane protein whose precise physiological function remains incompletely defined^7^. While CMT1A and HNPP are classified as primary myelin disorders, the prevailing pathogenic model emphasizes secondary axonal degeneration as the driver of clinical deficits^8,9^. However, despite severe motor impairments, reduced compound muscle action potentials, and significant muscle atrophy observed in CMT1A mouse models^10,11^, we and others have reported no clear evidence of denervation or axonal loss^10,12^. These findings implicate primary myelin dysfunction as a central contributor to disease pathogenesis and highlight the need to better understand the structural and molecular consequences of PMP22 dysregulation in myelin.

PMP22 is structurally related to Claudins, a tight junction protein family (pfam00822), and shares a similar transmembrane topology despite having only 25% sequence identity and 41% sequence similarity with Claudin-1. Upon comparing AlphaFold-predicted structures, we observed notable similarity between the extracellular loops of PMP22 and Claudin-1, suggesting a potentially conserved role in mediating adhesion.

Supporting this idea, PMP22 has been shown to localize to cell junctions in epithelial and endothelial cells, where its overexpression increases transepithelial electrical resistance^13–15^. In peripheral nerve, PMP22 is localized to compact myelin and has been linked to the organization of both tight junctions and adherens junctions. Prior studies have shown that key junctional components, including Claudin-19, ZO-1, ZO-2, JAM-C, and E-Cadherin, are disrupted in PMP22-deficient mice^16–18^. In HNPP model mice (*PMP22*+/–), Claudin-19 shows reduced and asymmetric accumulation near Nodes of Ranvier, while in *PMP22*–/– mice, E-Cadherin is abnormally dispersed along internodes rather than enriched at Schmidt-Lanterman incisures (SLIs)^16,18^. These findings further support an adhesion-related role for PMP22 in myelinating Schwann cells.

To further investigate whether PMP22 regulates adhesion and how *PMP22* gene dosage impacts myelin integrity in CMT1A and HNPP model mice, we systematically analyzed teased peripheral nerve fibers using high-resolution confocal imaging. Our data revealed that altered *PMP22* dosage leads to marked disruptions in adherens junction components, including E-Cadherin, β-Catenin, and F-Actin, which are prominently localized to SLIs. These disruptions were more pronounced in CMT1A, where SLIs appeared abnormally compacted relative to wildtype nerves. In contrast, nerves from HNPP model mice exhibited a more dispersed SLI architecture, particularly evident through F-Actin labeling, while E-Cadherin and β-Catenin showed more modest changes. These findings suggest that PMP22 dosage inversely affects adherens junction organization and likely SLI morphology. Importantly, not all SLI-resident proteins were affected in the same manner. We also examined additional adhesion-related proteins, including CADM4, Connexin29 (Cx29), and Myelin-Associated Glycoprotein (MAG). While CADM4 distribution remained unchanged in both CMT1A and HNPP, Cx29 and MAG displayed distinct alterations compared to adherens junction components. Specifically, both proteins exhibited more diffuse localization and increased signal intensity at SLIs in CMT1A and HNPP nerves, with changes generally more pronounced in CMT1A. These findings highlight selective and protein-specific molecular changes at SLIs that likely influence the structure and function of this specialized compartment.

Given the anatomical and functional link between SLIs and the Node of Ranvier via the inner mesaxon and prior studies showing that Cx29 associates with Kv1 channels at juxtaparanodes^19^, we next examined the molecular organization of Nodes of Ranvier. All three nodal domains were affected, with changes typically correlating with disease severity: CMT1A nerves showed more pronounced alterations than HNPP. At the node, voltage-gated sodium channels (Nav) exhibited evidence of nodal widening. At juxtaparanodes and paranodes, Kv1.2 and Caspr, respectively, demonstrated reduced enrichment and/or abnormal spreading along the internode. Patterning defects at the Nodes of Ranvier are likely to directly impair nerve conduction, whereas changes at the SLIs may have a more indirect effect, potentially by disrupting metabolic support. Therefore, our findings support a model in which PMP22 plays an essential role in organizing myelin architecture, likely through the regulation of adherens junctions. Disruption of this function is predicted to impair Schwann cell metabolic support and axonal ion homeostasis, thereby compromising both the structural integrity and functional capacity of the myelin sheath. This work challenges the long-standing axon-centric paradigm of CMT1A and HNPP pathogenesis and instead positions primary myelin dysfunction as a central driver of disease. Furthermore, it identifies junctional complexes as actionable molecular targets, providing mechanistic insight and a conceptual framework with potential relevance to other dysmyelinating peripheral neuropathies.

## Materials and Methods

### Animal Husbandry

All experiments were conducted with approval from the Johns Hopkins University and University of Missouri Animal Care and Use Committees. C3-PMP mice (B6.Cg-Tg(PMP22) C3Fbas/J, referred to as CMT1A mice) were obtained from the Jackson Laboratory (Stock #: 030052), maintained/expanded as heterozygotes by breeding with C57BL/6J wildtype mice (the Jackson Laboratory, Stock #: 000664) and genotyped according to the suggested protocol. LacZ PMP22 Deficient mice (referred to as HNPP mice) were obtained from Regeneron via Dr. Lucia Notterpek, maintained/expanded as heterozygotes by breeding with 129S1/SvImJ wildtype mice (the Jackson Laboratory, Stock #: 002448) and genotyped according to the suggested protocol^20^. A balanced representation of male and female mice aged 3 months were perfused and their sciatic nerves were harvested to prepare teased nerve fibers for immunostaining.

### Perfusion and Teased Nerve Fibers

Mice were anesthetized with Isoflurane and transcardiac perfusion was performed under anesthesia. The blood was cleared by perfusion with chilled 1x PBS followed by perfusion with chilled 4% paraformaldehyde prepared in 1x PBS. After perfusion, both sciatic nerves were dissected and post-fixed in 4% paraformaldehyde for three hours at 4° C. The nerves were then washed five times quickly with cold 1x PBS and stored at 4° C overnight. The next day the tibial branch was isolated in a dish of cold 1x PBS and cut into four segments. Each segment was transferred to a drop of 1x PBS on a Superfrost Plus microscope slide (Fisher Scientific) and gently teased with fine needles to separate the nerve fibers. After teasing, excess PBS was removed and the teased nerve fiber slides were dried at room temperature overnight. The slides were stored at -80° C the next day until ready for immunofluorescence staining.

### Immunofluorescence Staining of Teased Nerve Fibers

Teased nerve fibers for immunofluorescence were permeabilized by submerging the slides in -20° C chilled Acetone in Coplin jars for ten minutes while rocking at room temperature. After removing the acetone, the fibers were washed with 1x PBS three times for five minutes each while rocking at room temperature. The edges of each slide were quickly dried and a boundary around the fibers was drawn using a PAP pen. This was performed one slide at a time to prevent drying of the nerve fibers. Slides were resubmerged in the last 1x PBS wash after the PAP boundary was dry. Excess 1x PBS was removed from each slide before placing the slides into an immunostaining tray and adding 0.5ml of antibody buffer per slide (190ml – 1x PBS, 10ml - 40-50% Fish Gelatin [Sigma Aldrich, G7765] and 200ul Triton X-100, 10ml aliquots stored at -20° C). This was performed in batches of three slides to prevent drying of the nerve fibers. The nerve fibers were blocked with antibody buffer at room temperature for one hour. Primary antibody solutions (see Table 1 for concentrations) were prepared in antibody buffer on ice. After removing the antibody buffer used for blocking, 0.5ml of primary antibody solution was added to each slide and incubated at 4° C overnight.

**Table 1.** Primary Antibodies used in this Study.

| Primary Antibody | Company, Cat. Number | Species | Concentration |
| --- | --- | --- | --- |
| anti-E-Cadherin | Thermo Fisher, 14-3249-82 | Rat | 1:100 |
| anti-Neurofascin155 (Pan) | R&D Systems, AF3235 | Chicken | 1:500 |
| anti-ZO-1 | DSHB, R26.4C | Rat | 1:50 |
| anti-β-Catenin | Sigma-Aldrich, C2206 | Rabbit | 1:500 |
| anti-p120-Catenin | Thermo Fisher, PA5-82545 | Rabbit | 1:500 |
| anti-CADM4 | Thermo Fisher, MA5-24141 | Rat | 1:100 |
| anti-Connexin29 | Thermo Fisher, 34-4200 | Rabbit | 1:50 |
| anti-Nav (Pan) | Absolute Antibody, Ab02113-8.1 | Rat | 1:500 |
| anti-Kv1.2 | Absolute Antibody, Ab02104-23.0 | Rabbit | 1:500 |
| anti-Caspr1 | Abcam, ab34151 | Rabbit | 1:1000 |

The nerve fibers were washed in 1x PBS three times for ten minutes each while rocking at room temperature. Secondary antibody solutions (see Table 2 for concentrations) were prepared in antibody buffer on ice. Excess 1x PBS was removed from each slide before placing the slides into an immunostaining tray and adding 0.5ml of secondary antibody solution to each. This was performed in batches of three slides to prevent drying of the nerve fibers. The nerve fibers were incubated with secondary antibody at room temperature for one and a half hours.

**Table 2.** Secondary and Direct Conjugated Antibodies used in this Study.

| Secondary or Direct Conjugated Antibody | Company | Concentration |
| --- | --- | --- |
| 405 Phalloidin | Thermo Fisher, A30104 | 1:400 |
| MAG Alexa Fluor 488 | Sigma-Aldrich, MAB1567A4 | 1:100 |
| Goat anti-Rabbit Alexa Fluor 488 | Thermo Fisher, A11034 | 1:1000 |
| Goat anti-Rat Alexa Fluor 568 | Thermo Fisher, A11077 | 1:1000 |
| Goat anti-Chicken Alex Fluor 647 | Thermo Fisher, A11041 | 1:1000 |

The nerve fibers were washed in 1x PBS three times for ten minutes each while rocking at room temperature. Excess 1x PBS was removed, 60μl Prolong Gold antifade reagent (Thermo Fisher, P36930) was added to each and topped with No. 1.5H Precision cover glass. Slides were dried at room temperature overnight in the dark and stored at -20° C the next day until ready for imaging.

### Confocal Immunofluorescence Imaging, Data Analysis and Statistics

Confocal imaging was performed using a Zeiss LSM800, Zeiss LSM880 or Olympus FV4000 using a 60x oil objective. Z-stacks were acquired from at least six different regions per mouse per condition and analyzed using Imaris x64 version 9.2.1 (Oxford Instruments, Bitplane). Five SLIs from at least three different nerve fibers were manually outlined in each image to create isosurfaces. Mean signal intensity was measured within each isosurface as a readout of SLI mean signal intensity. The signal for a channel of interest was then duplicated only within each SLI isosurface and a threshold was established as real signal for each antibody. A second isosurface was created within each SLI isosurface by setting the established signal threshold for each antibody. The volume of the second isosurface and the diameter of the nerve fiber at each SLI (axon + myelin sheath) were recorded. The distribution at SLIs was calculated by normalizing each thresholded signal volume to the fiber diameter for that SLI. Nodes of Ranvier, generally from two to three different nerve fibers, were analyzed similarly by manually tracing a node and the adjacent internodal myelin sheaths on both sides of the node up until the first SLI or a cross point with another nearby myelinated fiber. Mean signal intensity was measured within each node + proximal internode isosurface as a readout of Node of Ranvier mean signal intensity. The signal for a channel of interest was then duplicated only within each node + proximal internode isosurface and a threshold was established as real signal for each antibody. A second isosurface was created within each node + proximal internode isosurface by setting the established signal threshold for each antibody. The volume of the second isosurface and the diameter of the nerve fiber (axon + myelin sheath) were recorded. The distribution at Nodes of Ranvier was calculated by normalizing each thresholded signal volume to the fiber diameter. Internodal spread for Kv1.2 and Caspr was measured by selecting only the second isosurfaces near the node using the circle selection tool in Imaris and subtracting the volume at the node from the total volume and normalizing this to the total node + proximal internode isosurface volume.

All data were evaluated for significance by t-test using Prism 9 (GraphPad). For each experiment, the data were analyzed by three statistical tests: (1) unpaired t-test on all individual datapoints, (2) unpaired t-test on the average datapoints from each biological replicate and (3) nested t-test on all datapoints grouped by biological replicate. Significance detected (p < 0.05) with all three t-test statistics is denoted as ***, with only two of these t-test statistics is denoted at ** and with only one is denoted as *. The statistical test used, exact n, definition of what n represents, and graphical display details are located in the figure legends. Randomization was performed for all experiments by randomly selecting mice from multiple litters and representing both sexes in each cohort. No data points from any assay were excluded from this study.

## Results

### Altered Adherens Junction Organization at Schmidt-Lanterman Incisures in CMT1A and HNPP

PMP22 belongs to the Claudin superfamily, a group of transmembrane proteins best known for forming tight junctions that regulate paracellular permeability and maintain selective barriers. Consistent with this, PMP22 shares a homologous protein topology, and its extracellular loops exhibit striking structural similarity to the orthodox Claudins, like Claudin-1, in both humans and mice (Figure 1a,b and Supplemental Figure 1a). In contrast, ClustalW sequence alignments reveal only modest sequence similarity between PMP22 and Claudin-1, despite strong conservation of PMP22 across species (Supplemental Figure 1b). These findings reinforce the notion that PMP22 may carry out Claudin-like functions through conserved structural features rather than high primary sequence identity, particularly in the context of adhesion junctions in peripheral nerve myelin. In myelinating Schwann cells, adhesion junctions including tight and adherens junctions, localize to specialized cytoplasmic-like domains including SLIs and Nodes of Ranvier, where they are believed to play essential roles in structural integrity and function^21–23^ (Figure 1c).

**Figure 1.**
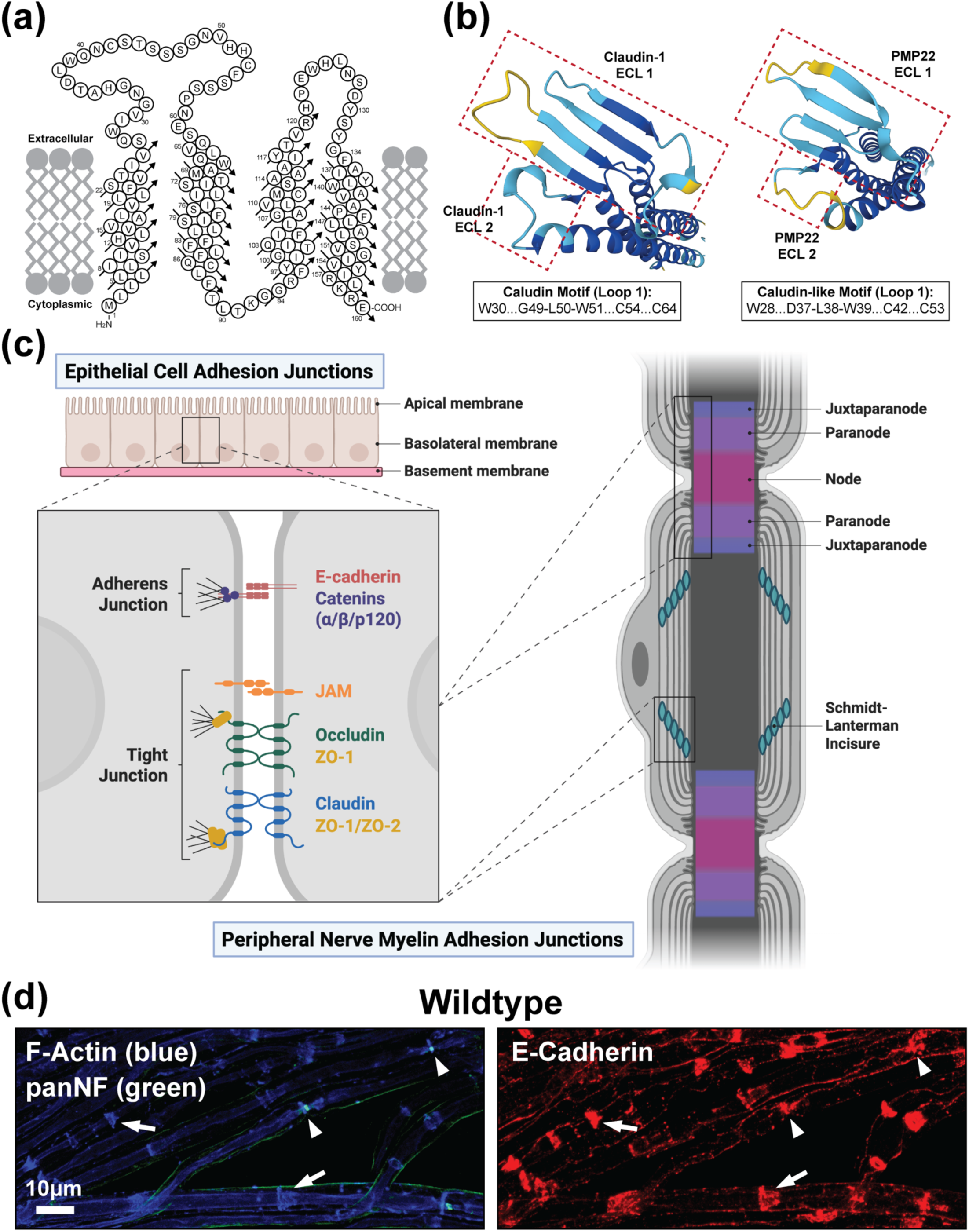
Support for PMP22 Function in Cell Adhesion and Peripheral Nerve Myelin Adhesion Junctions. **(a)** PMP22 is a member of the Claudin superfamily of proteins. Orthodox Claudins establish selective barriers as components of tight junctions and as such PMP22 protein topology is homologous. Topology for human PMP22, adapted from^34^. **(b)** AlphaFold predicted structures of human Claudin-1 and human PMP22 extracellular loops (ECLs) viewed from above (AlphaFold Protein Structure Database^35^). The structural similarity between the ECLs of human Claudin-1 and human PMP22 is striking suggesting that PMP22 functions similarly to Claudin proteins. **(c)** Adhesion junctions, including adherens junctions and tight junctions, are localized to specialized cytoplasmic-like compartments in peripheral nerve myelin including Schmidt-Lanterman Incisures (SLIs) and Nodes of Ranvier. Created in https://BioRender.com. **(d)** 3-month-old wildtype (WT, C57BL/6J) teased tibial nerve fibers stained for the adherens junction protein E-Cadherin (red) showing prominent localization at SLIs (arrows), Nodes of Ranvier (arrowheads) and inner mesaxons. Scale bar, 10μm.

To investigate the potential role of PMP22 in adhesion, we examined peripheral nerve myelin from constitutive mouse models of CMT1A (C3-PMP)^11^ and HNPP (PMP22-deficient LacZ)^20^. Teased nerve fibers were prepared from adult mice (aged three months) by dissecting sciatic nerves, isolating the tibial branches and performing confocal immunofluorescence imaging. The tight junction component ZO-1 demonstrated localization at Nodes of Ranvier (data not shown) and the adherens junction component E-Cadherin was prominently localized to SLIs, inner mesaxons and at Nodes of Ranvier (Figure 1d). CMT1A fibers did not show marked changes in ZO-1 expression or localization (data not shown); however, E-Cadherin was severely disrupted, with striking disorganization particularly evident at SLIs (Figure 2a and Supplemental Figure 2a).

**Figure 2.**
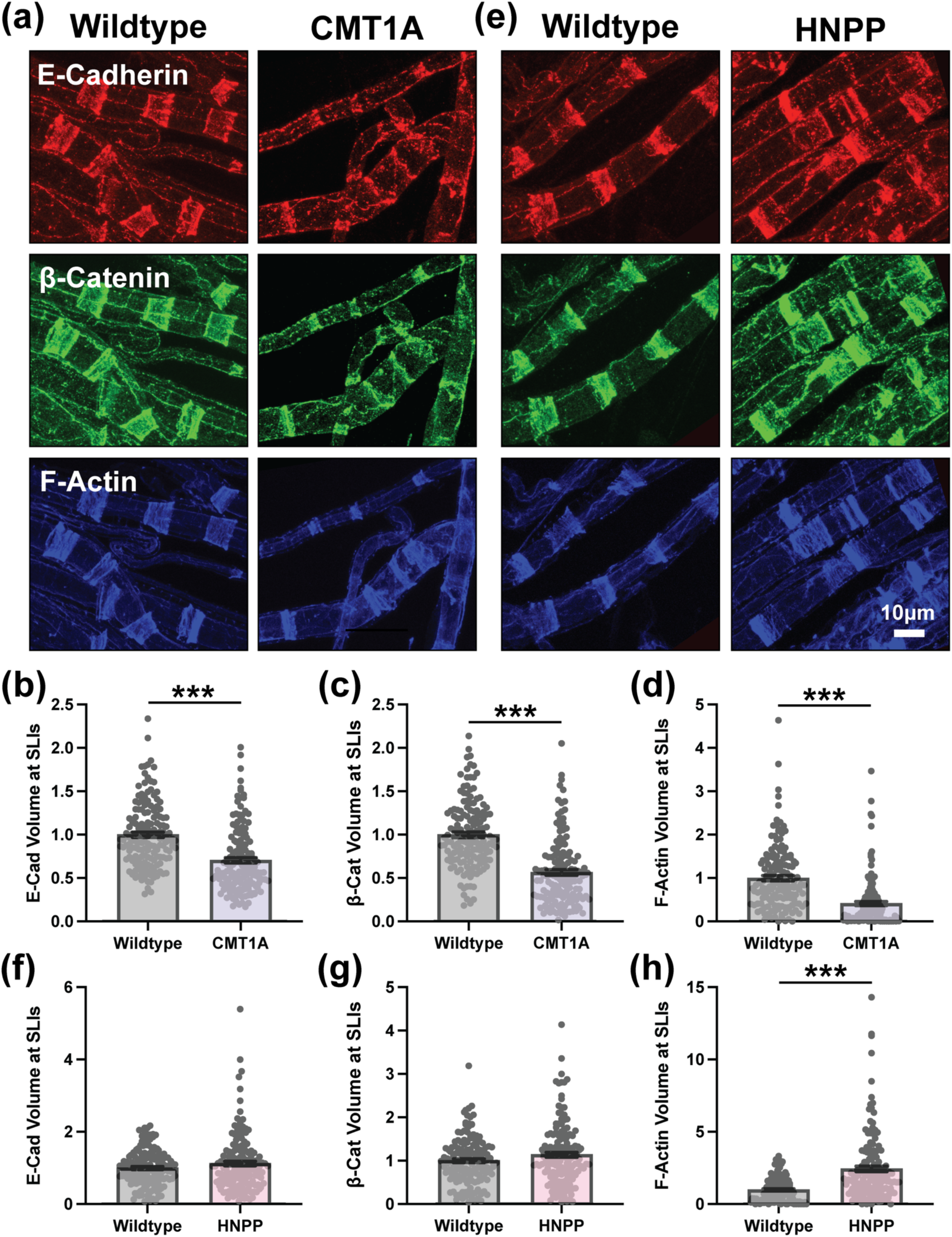
Altered Adherens Junction and F-Actin Patterning at SLIs in CMT1A and HNPP Model Peripheral Nerve Myelin. Representative images of 3-month-old **(a)** wildtype (WT, C57BL/6J) and CMT1A or **(e)** WT (129S1/SvImJ) and HNPP teased tibial nerve fibers stained for the adherens junction complex proteins E-Cadherin (red), β-Catenin (green) and F-Actin (blue). Quantification of protein distribution at SLIs (signal volume/fiber diameter) for **(b, f)** E-Cadherin, **(c, g)** β-Catenin and **(d, h)** F-Actin in CMT1A and HNPP, respectively. n=5 animals (∼30 SLIs/animal). Bar graphs represent mean ± SEM with individual data points. ***p<0.05 demonstrate statistical significance with three separate t-test statistics (unpaired t-test with all individual data points, unpaired t-test with experimental average data points and nested t-test). Scale bar, 10μm.

The signal is more punctate and the funnel shape structure of SLIs is often disrupted. Quantification of this defect revealed reduced E-Cadherin signal volume as normalized to fiber diameter (here on referred to as distribution), to account for the loss of large caliber myelinated fibers in CMT1A model mice, or a compacted distribution (Figure 2b; 0.704-fold change, nested t-test p=0.0118). Mean E-Cadherin signal intensity at SLIs was modestly increased reaching statistical significance but is likely not biologically significant due to the small fold change (Supplemental Figure 2b; 1.098-fold change, unpaired t-test with individual data points p=0.0007). The core complex of adherens junctions consists of the single pass transmembrane protein E-cadherin and three cytoplasmic Catenin proteins (p120-Catenin, β-Catenin and α-Catenin) which connect the cadherin complex to the actin cytoskeleton and multiple signaling pathways^24^. The distributions of β-Catenin and F-Actin mimic the changes in E-Cadherin but exhibit even stronger deficits (Figure 2a,c,d; 0.567-fold change, nested t-test p=0.0028 and 0.414-fold change, nested t-test p=0.0024, respectively). Whereas mean β-Catenin and F-Actin signal intensity at SLIs is unchanged in CMT1A (Supplemental Figure 2a,c,d). These findings suggest that adherens junctions at SLIs and likely the overall morphology SLIs, as evidenced by dramatic changes in F-actin, are compromised in CMT1A model myelin demonstrating an abnormally compacted SLI pattern.

In HNPP model myelin, these adherens junction components remained largely unchanged, with only mean β-Catenin signal intensity at SLIs reaching statistical significance (1.106-fold change, unpaired t-test with individual data points p=0.0033) but likely lacking biological relevance (Figure 2e–g and Supplemental Figure 2e–g). However, SLI architecture appears oppositely affected in HNPP as evidenced by a more dispersed distribution and an increased mean signal intensity of F-actin (Figure 2e,h and Supplemental Figure 2e,h; 2.468- fold change in distribution, nested t-test p=0.0044 and 1.352-fold change in mean signal intensity, nested t-test p=0.0316). β-Catenin and F-actin are key components of adherens junctions, where they contribute to both structural stability and dynamic regulation of cell adhesion. β-Catenin binds the intracellular domain of E-Cadherin and associates with α-Catenin, which in turn links to F-Actin. Together, these interactions bridge adherens junctions with the actin cytoskeleton, supporting adherens junction integrity. In peripheral nerve, α3- Catenin is the predominant isoform for which reliable antibodies are limited, preventing its inclusion in this study^25^. p120-Catenin also binds to the intracellular domain of E-Cadherin serving to regulate E-Cadherin stability by blocking endocytosis and turnover as well as modulating small GTPase activity to influence actin dynamics and junction organization. p120-Catenin distribution at SLIs was unchanged in myelin from CMT1A model mice but was significantly increased in HNPP (Figure 3a-d and Supplemental Figure 3a; 2.002-fold change, unpaired t-test with individual data points p<0.0001). However, p120-Catenin mean signal intensity at SLIs was modestly increased in both CMT1A (1.091-fold change, unpaired t-test with individual data points p=0.0400) and HNPP (1.112-fold change, unpaired t-test with individual data points p=0.0033) although unlikely biologically significant (Figure 3a,c and Supplemental Figure 3a-d). Given that p120-Catenin stabilizes E-Cadherin at the membrane^26^, its increased expression or localization at SLIs may represent a compensatory response to adherens junction disruption, potentially resulting in better preservation of junctional integrity in HNPP than CMT1A.

**Figure 3.**
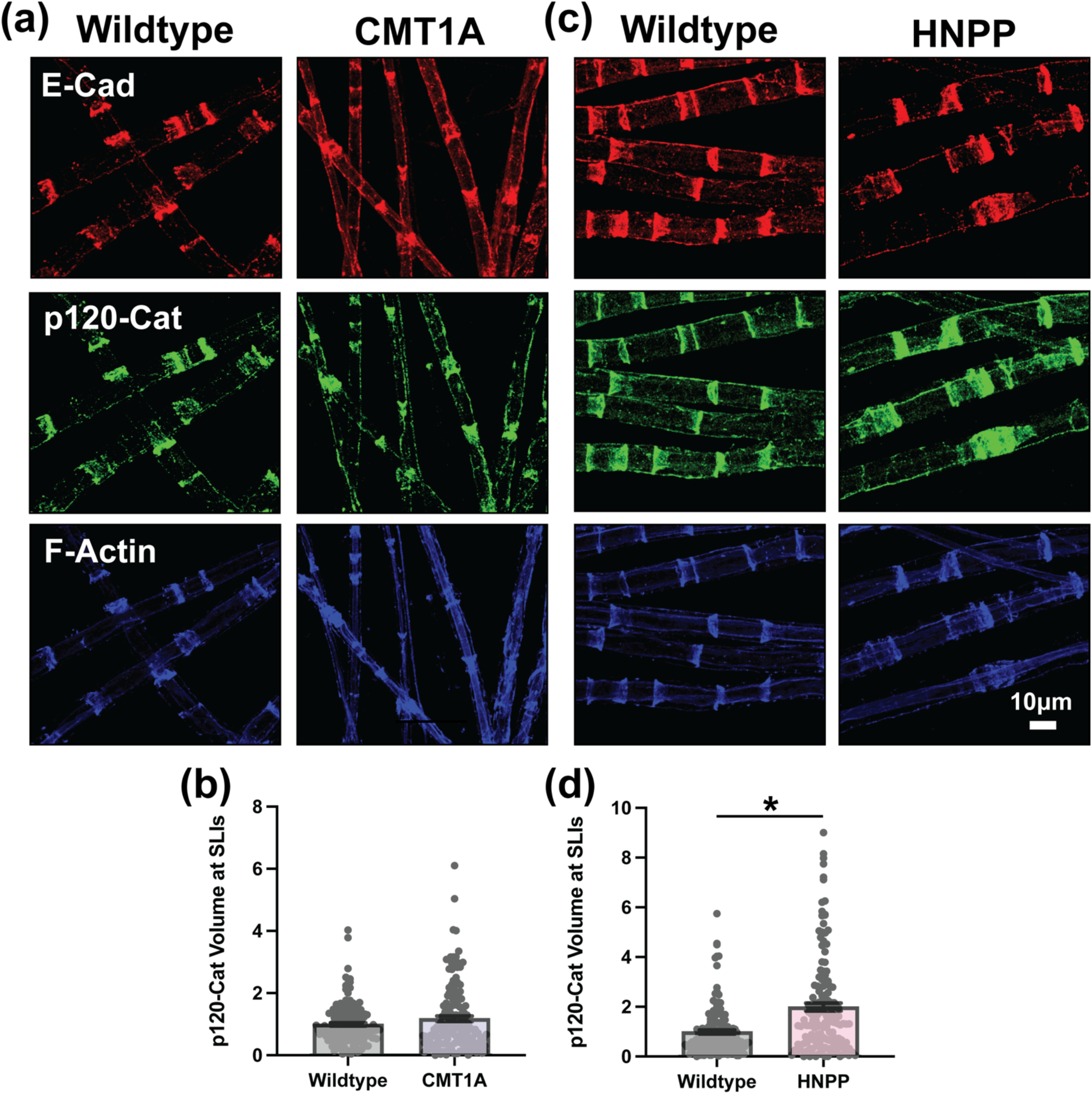
Variable Disruption of p120-Catenin at SLIs in CMT1A and HNPP Model Myelin. Representative images of 3-month-old **(a)** WT (C57BL/6J) and CMT1A or **(c)** WT (129S1/SvImJ) and HNPP teased tibial nerve fibers stained for the adherens junction complex proteins E-Cadherin (red), p120-Catenin (green) and F-Actin (blue). Quantification of p120-Catenin protein distribution at SLIs (signal volume/fiber diameter) in **(b)** CMT1A and **(d)** HNPP. n=5 animals (∼30 SLIs/animal). Bar graphs represent mean ± SEM with individual data points. *p<0.05 demonstrate statistical significance with one t-test statistic (unpaired t-test with all individual data points). Scale bar, 10μm.

### Additional Schmidt-Lanterman Incisures Abnormalities with Divergent Features in CMT1A versus HNPP

Adherens junctions function to maintain structural integrity^27^ and F-Actin is commonly used as a marker of SLIs suggesting that gross SLI morphology is disrupted in CMT1A and HNPP. Therefore, we next determined if multiple other SLI components mimicked the effects observed with adherens junction components and F-Actin. Cell Adhesion Molecule 4 (CADM4 but also known as SynCAM4 or Necl-4), Connexin29 (Cx29), and Myelin-Associated Glycoprotein (MAG) are key molecular components of peripheral nerve myelin that are enriched at SLIs and contribute to axoglial communication and structural organization^19,28–30^. CADM4 and MAG are adhesion-like proteins and Cx29 is a gap junction protein. CADM4 mean signal intensity and distribution at SLIs in unchanged in both CMT1A and HNPP indicating that not are SLI components are disrupted (Figure 4a,b,e,f and Supplemental Figure 4a-d). However, Cx29 and MAG show striking alterations in their distribution and mean signal intensity at SLIs in CMT1A and HNPP, differing from the pattern observed for F-actin and adherens junctions. Cx29 distribution was dramatically more dispersed at SLIs in CMT1A (Figure 4a,c and Supplemental Figure 4a; 6.233-fold change, unpaired t-test with individual data points p<0.0001) and more modestly in HNPP (Figure 4e,g and Supplemental Figure 4c; 1.257-fold change, unpaired t-test with individual data points p=0.0205). Similarly, Cx29 mean signal intensity at SLIs was increased more so in CMT1A (Figure 4a,d and Supplemental Figure 4a; 1.596-fold change, nested t-test p=0.0162) than HNPP (Figure 4e,h and Supplemental Figure 4c; 1.132-fold change, unpaired t-test with individual data points p=0.0205).

**Figure 4.**
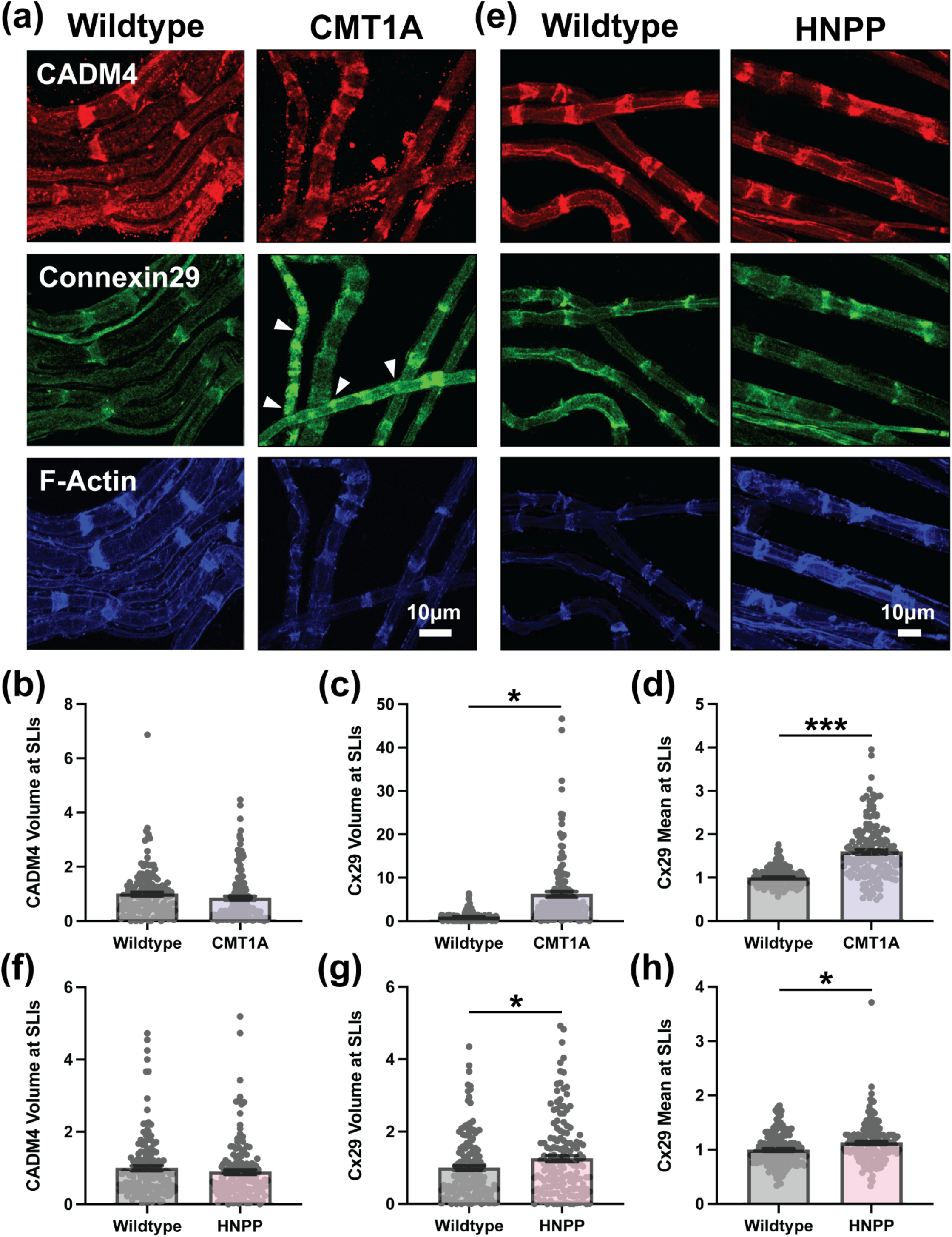
Distinct Connexin29 Patterning Defects but Unchanged CADM4 at SLIs in Peripheral Nerve Myelin from CMT1A and HNPP Model Mice. Representative images of 3-month-old **(a)** WT (C57BL/6J) and CMT1A or **(e)** WT (129S1/SvImJ) and HNPP teased tibial nerve fibers stained for CADM4 (red), Connexin29 (Cx29, green) and F-Actin (blue). Note the focal accumulations of Cx29 outside the SLI compartment in CMT1A model myelin (arrowheads). Quantification of protein distribution at SLIs (signal volume/fiber diameter) for **(b, f)** CADM4 and **(c, g)** Cx29 in CMT1A and HNPP, respectively. Quantification of mean Cx29 signal intensity at SLIs (readout of protein expression levels/localization) in **(d)** CMT1A and **(h)** HNPP. n=5 animals (∼30 SLIs/animal). Bar graphs represent mean ± SEM with individual data points. ***p<0.05 demonstrate statistical significance with three separate t-test statistics (unpaired t-test with all individual data points, unpaired t-test with experimental average data points and nested t-test). *p<0.05 demonstrate statistical significance with one t-test statistic (unpaired t-test with all individual data points). Scale bars, 10μm.

MAG distribution at SLIs is increased in CMT1A (2.232-fold change, unpaired t-test with individual data points p<0.0001) and HNPP (2.471-fold change, unpaired t-test with individual data points p=0.0002) to a similar degree (Figure 5a,b,d,e). MAG mean signal intensity at SLIs is also comparably affected in CMT1A (Figure 5a,c; 1.290-fold change, unpaired t-test with individual data points p<0.0001) and HNPP (Figure 5d,f; 1.270-fold change, unpaired t-test with individual data points p<0.0001). These results collectively reveal that not all SLI components are equally affected in CMT and HNPP. Particularly notable are components that show opposite effects in CMT1A and HNPP, reflecting the inverse changes in *PMP22* copy number, such as F-Actin distribution. Another interesting pattern involves features more severely disrupted in CMT1A than in HNPP, aligning with the relative severity of patient symptoms, such as Cx29 distribution and mean signal intensity.

**Figure 5.**
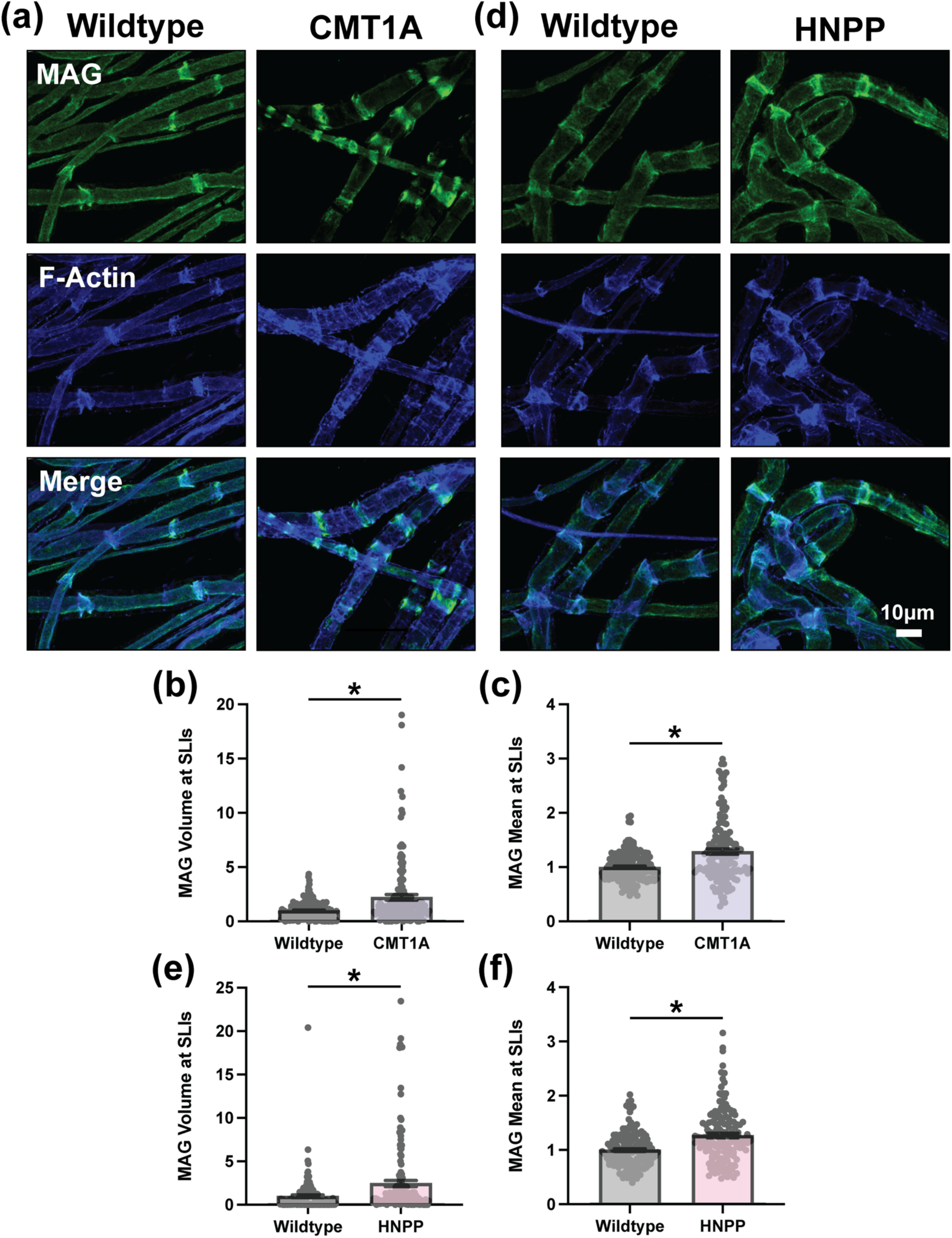
MAG Exhibits Comparable Mispatterning to Connexin29 at SLIs in CMT1A and HNPP Model Myelin. Representative images of 3-month-old **(a)** WT (C57BL/6J) and CMT1A or **(d)** WT (129S1/SvImJ) and HNPP teased tibial nerve fibers stained for MAG (green) and F-Actin (blue). Quantification of **(b, e)** MAG protein distribution (signal volume/fiber diameter) and **(c, f)** MAG mean signal intensity (readout of protein expression levels/localization) at SLIs in CMT1A and HNPP, respectively. n=5 animals (∼30 SLIs/animal). Bar graphs represent mean ± SEM with individual data points. *p<0.05 demonstrate statistical significance with one t-test statistic (unpaired t-test with all individual data points). Scale bar, 10μm.

Cx29 is also frequently observed as focal accumulations along the internode outside of SLIs (Figure 4a and Supplemental Figure 4a), suggesting that additional mechanisms beyond SLI-specific disruptions may occur in CMT1A and HNPP. Given that Cx29 and other SLI-associated proteins also localize to the inner mesaxon and near the Node of Ranvier, we turned our attention to nodal architecture, both due to its critical role in action potential propagation and because nodopathy is a well-established mechanism in acquired neuropathies.

### Peripheral Nerve Ion Channel Deficits at Nodes of Ranvier in CMT1A and HNPP

Nodes of Ranvier are spatially organized into three distinct regions, the node, paranode and juxtaparanode, to allow for saltatory conduction of action potentials along axons (Figure 6a). The node contains voltage-gated sodium channels critical for depolarization, with Nav1.6 predominating at mature nodes that support high-frequency firing, while Nav1.7, Nav1.8, and Nav1.9 are primarily enriched at nodes in sensory neurons^31,32^. The juxtaparanode contains voltage-gated potassium channels essential for repolarization, with Kv1.1 and Kv1.2 as the predominant subtypes in fast-conducting peripheral nerve axons^19^. And the paranode contains septate-like junctions formed by axonal Caspr and Contactin and Schwann cell Neurofascin 155, which anchor the terminal loops of each myelin wrap to the axon and functionally separate the node from the juxtaparanode^33^. Further motivation to investigate Node of Ranvier architecture arises from findings that Cx29 co-localizes with Kv1 channels at juxtaparanodes, suggesting electrical coupling between axons and myelinating Schwann cells, and that adherens junctions localize to paranodes where they are proposed to contribute to paranodal integrity^19,21^. Nodes and juxtaparanodes were assessed in teased nerved fibers from adult CMT1A and HNPP model mice by staining with anti-pan Nav and anti-Kv1.2 antibodies, respectively.

**Figure 6.**
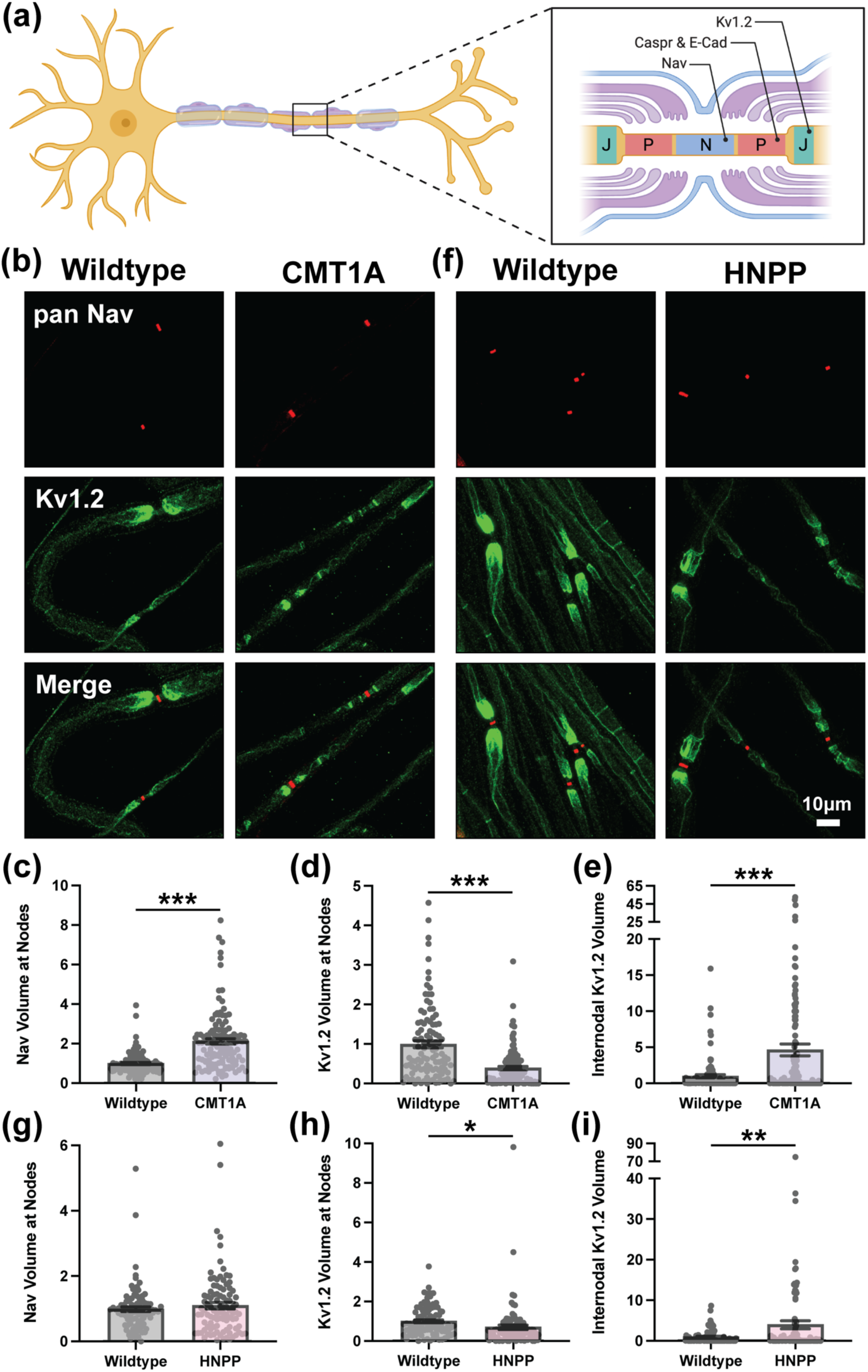
Disrupted Ion Channel Organization at Nodes of Ranvier in CMT1A and HNPP Model Peripheral Nerve Myelin. **(a)** Diagram demonstrating the molecular organization of three distinct domains at Nodes of Ranvier: nodes (N, containing Nav), paranodes (P, containing Caspr and E-Cadherin) and Juxtaparanodes (J, containing Kv1.2). Created in https://BioRender.com. Representative images of 3-month-old **(b)** WT (C57BL/6J) and CMT1A or **(f)** WT (129S1/SvImJ) and HNPP teased tibial nerve fibers stained with anti-Pan Nav (red) and anti-Kv1.2 (green). Quantification of protein distribution at Nodes of Ranvier (signal volume/fiber diameter) for **(c, g)** Nav and **(d, h)** Kv1.2 in CMT1A and HNPP, respectively. **(e, i)** Quantification Kv1.2 internodal spread away from Nodes of Ranvier in CMT1A and HNPP, respectively. n=5 animals (∼15-25 nodes/animal). Bar graphs represent mean ± SEM with individual data points. ***p<0.05 demonstrate statistical significance with three separate t-test statistics (unpaired t-test with all individual data points, unpaired t-test with experimental average data points and nested t-test). **p<0.05 demonstrate statistical significance with two separate t-test statistics (unpaired t-test with all individual data points and unpaired t-test with experimental average data points). *p<0.05 demonstrate statistical significance with one t-test statistic (unpaired t-test with all individual data points). Scale bar, 10μm.

Nav distribution at nodes was increased in CMT1A (2.063-fold change, nested t-test p=0.0125) but unchanged in HNPP (Figure 6b,c,f,g and Supplemental Figure 5a,d). Additionally, Nav mean signal intensity at nodes was increased in CMT1A (1.570-fold change, unpaired t-test with individual data points p<0.0001) but unchanged in HNPP (Figure 6b,f and Supplemental Figure 5a,b,d,e). Together, these results indicate that nodal widening is prominent in CMT1A model myelin but not in HNPP.

We also observed pronounced alterations in Kv1.2 patterning in CMT1A and HNPP model nerves. Kv1.2 enrichment at the juxtaparanodes was frequently reduced, with a more pronounced distribution reduction in CMT1A (Figure 6b,d and Supplemental Figure 5a; 0.398-fold, nested t-test p=0.0002) as compared to HNPP (Figure 6f,h and Supplemental Figure 5d; 0.702-fold, unpaired t-test with individual data points p=0.0305). Kv1.2 mean signal intensity at and near Nodes of Ranvier was also reduced in CMT1A and HNPP but to a similar degree (Figure 6b and Supplemental Figure 5a,c; 0.800-fold change in CMT1A, unpaired t-test with individual data points p<0.0001 and Figure 6f and Supplemental Figure 5d,f; 0.792-fold change in HNPP, unpaired t-test with individual data points p<0.0001). In addition, Kv1.2 was frequently found to be spread along the internode away from the juxtaparanodal compartment. This was quantified by measuring Kv1.2 distribution along the proximal internode, calculated by subtracting the nodal Kv1.2 volume from the total Kv1.2 volume and normalizing to the total volume of the selected fiber segment. Internodal Kv1.2 distribution was markedly increased in both CMT1A and HNPP models, with a more pronounced elevation in CMT1A (Figure 6b,e and Supplemental Figure 5a; 4.769-fold change in CMT1A, nested t-test p=0.0205 and Figure 6f,i and Supplemental Figure 5d; 3.987-fold change in HNPP, unpaired t-test with individual data points p=0.0041). These robust disruptions in Node of Ranvier architecture, including nodal widening, reduced Kv1.2 juxtaparanodal enrichment and internodal Kv1.2 mislocalization, are likely to result in impaired action potential propagation, with more severe consequences in CMT1A than in HNPP.

### Compromised Paranode Integrity at Nodes of Ranvier in CMT1A and HNPP

Given the role of paranodal septate-like junctions in organizing Nodes of Ranvier, a key component of this complex, Caspr (also called Caspr1) was evaluated by confocal immunofluorescence imaging. Caspr plaques at the paranode were often fragmented and focal accumulations of Caspr were often visible along the internode similar to Kv1.2 spread. However, these abnormalities were comparable in magnitude between CMT1A and HNPP. Caspr distribution was increased (Figure 7a,c and Supplemental Figure 6a; 1.305-fold change in CMT1A, unpaired t-test with individual data points p=0.0002 and Figure 7e,g and Supplemental Figure 6d; 1.739-fold change in HNPP, unpaired t-test with individual data points p<0.0001) and Caspr mean signal intensity was modestly increased (Figure 7a and Supplemental Figure 6a,c; 1.106-fold change in CMT1A, unpaired t-test with individual data points p=0.0129 and Figure 7e and Supplemental Figure 6d,f; 1.255-fold change in HNPP, unpaired t-test with individual data points p<0.0001). Internodal Caspr distribution was quantified using the same method as for Kv1.2, revealing a dramatic increase in both CMT1A and HNPP, with a potentially greater extent in CMT1A (Figure 7a,d and Supplemental Figure 6a; 4.263-fold change in CMT1A, unpaired t-test with individual data points p=0.0034 and Figure 7e,h and Supplemental Figure 6d; 3.871-fold change in HNPP, unpaired t-test with individual data points p=0.0046).

**Figure 7.**
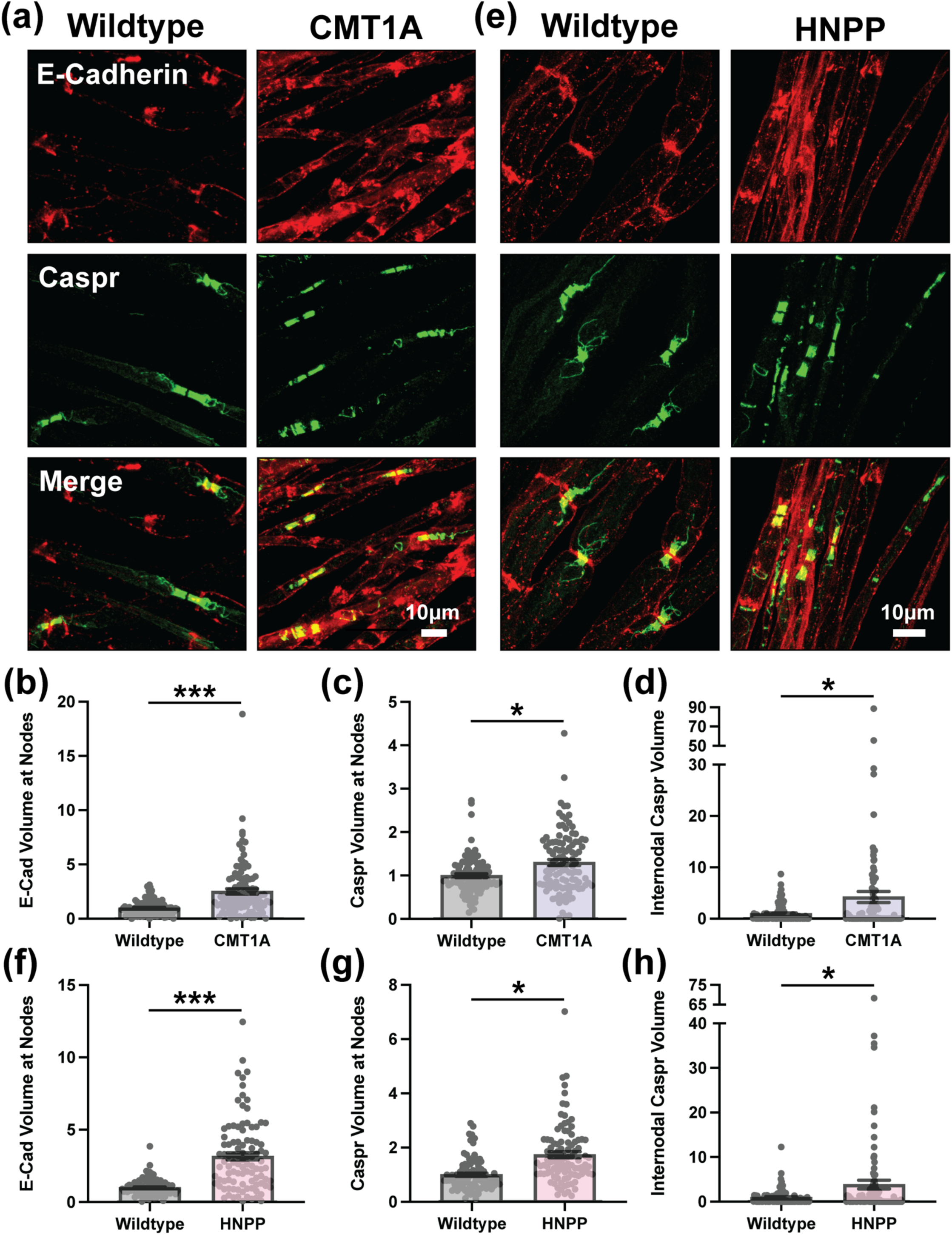
CMT1A and HNPP Model Myelin also Demonstrate Disruption of Caspr and E-Cadherin at Paranodes. Representative images of 3-month-old **(a)** WT (C57BL/6J) and CMT1A or **(e)** WT (129S1/SvImJ) and HNPP teased tibial nerve fibers stained for E-Cadherin (red) and Caspr (green). Quantification of protein distribution at Nodes of Ranvier (signal volume/fiber diameter) for **(b, f)** E-Cadherin and **(c, g)** Caspr in CMT1A and HNPP, respectively. **(e, i)** Quantification Caspr internodal spread away from Nodes of Ranvier in CMT1A and HNPP, respectively. n=5 animals (∼15-25 nodes/animal). Bar graphs represent mean ± SEM with individual data points. ***p<0.05 demonstrate statistical significance with three separate t-test statistics (unpaired t-test with all individual data points, unpaired t-test with experimental average data points and nested t-test). *p<0.05 demonstrate statistical significance with one t-test statistic (unpaired t-test with all individual data points). Scale bars, 10μm.

E-Cadherin localization at Nodes of Ranvier was also examined, given the striking alterations in adherens junction components at SLIs in CMT1A model myelin. E-Cadherin distribution was increased in both CMT1A and HNPP, with a potentially greater extent in HNPP (Figure 7a,b and Supplemental Figure 6a; 2.623- fold change in CMT1A, nested t-test p=0.0335 and Figure 7e,f and Supplemental Figure 6d; 3.118-fold change in HNPP, nested t-test p=0.0138), while mean signal intensity was similarly elevated in both models (Figure 7a and Supplemental Figure 6a,b; 1.324-fold change in CMT1A, nested t-test p=0.0061 and Figure 7e and Supplemental Figure 6d,e; 1.338-fold change in HNPP, nested t-test p=0.0239). The more pronounced alterations in Caspr and E-Cadherin distribution and mean signal intensity at the Node of Ranvier observed in HNPP, compared to CMT1A, do not correlate with the relative severity of patient symptoms. These findings are likely confounded by the presence of tomacula, focal myelin thickenings that frequently occur near the node in HNPP, complicating interpretation. This obscures the conclusions drawn from these experiments but lends greater weight to deficits more severely affected in CMT1A than in HNPP as potential primary drivers of disease pathogenesis (summarized in Figure 8).

**Figure 8.**
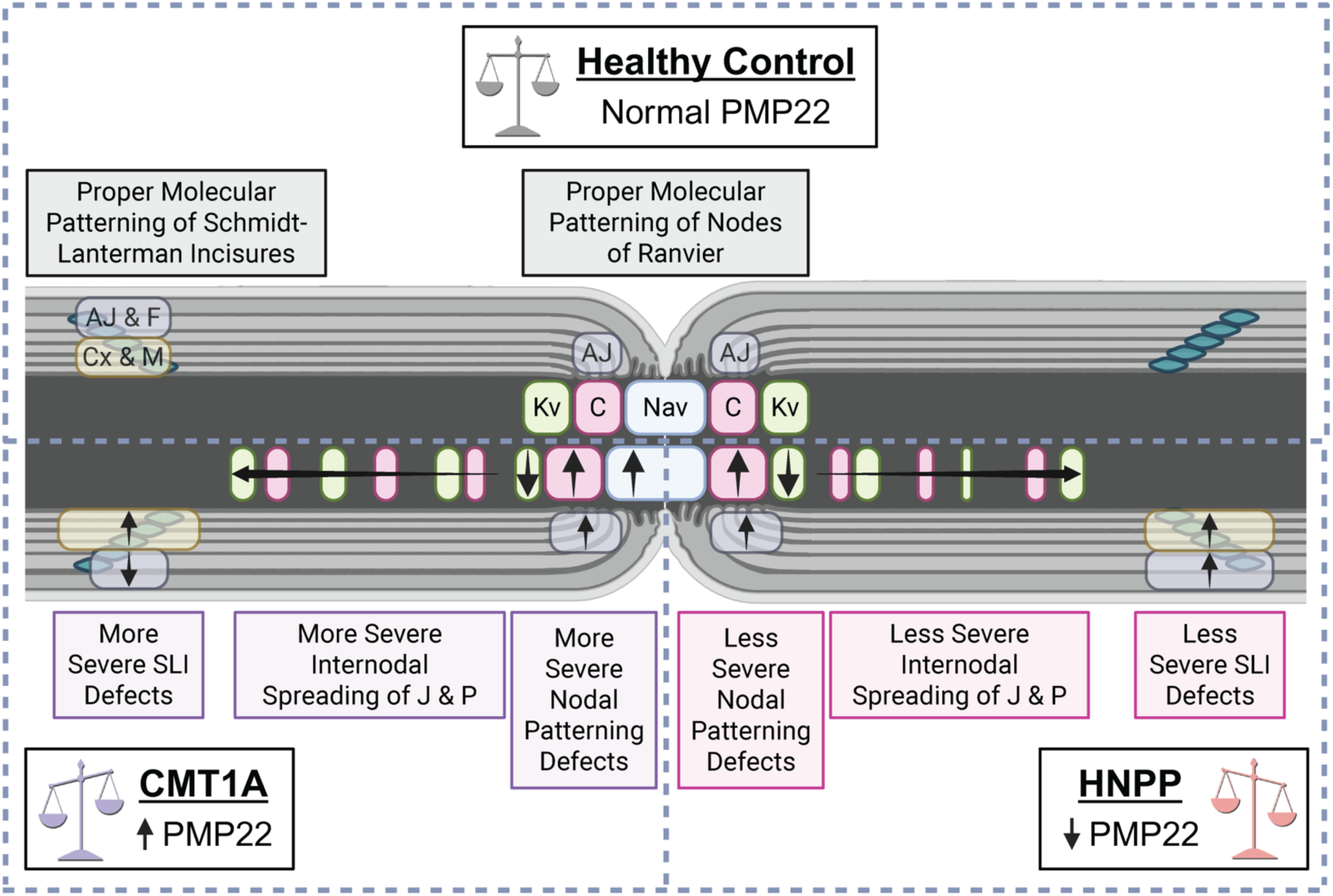
Overview of Molecular Alterations in Relation to CMT1A and HNPP Disease Mechanisms. Given that CMT1A patients typically exhibit more severe symptoms than HNPP patients, molecular or cellular phenotypes that are more pronounced in CMT1A, or show opposite patterns between the two, are likely to reflect key contributors to disease pathogenesis. Several of our observations that we report here fit this trend. In SLIs, E-Cadherin, β-Catenin and F-Actin demonstrate compacted distributions in CMT1A but are increased or unchanged in HNPP and Connexin29 is dramatically expanded in CMT1A and less so in HNPP. At Nodes of Ranvier, Kv1.2 is less concentrated at juxtaparanodes and is spread internodally in CMT1A more so than in HNPP, Nav demonstrates nodal widening in CMT1A but is unchanged in HNPP and Caspr demonstrates more internodal spread in CMT1A than in HNPP. These results suggest that changes at Nodes of Ranvier, such as nodal widening and disrupted domain-specific molecular patterning, along with accompanying abnormalities at SLIs, likely impair peripheral nerve myelin function, with more severe defects observed in CMT1A than HNPP. These structural disruptions may contribute to disease pathogenesis by impairing metabolic support and axonal ion homeostasis, resulting in slowed nerve conduction or conduction block despite grossly normal compact myelin. Created in https://BioRender.com.

## Discussion

Although secondary axonal degeneration has historically been proposed as the primary driver of functional deficits in CMT1A^8,9^, recent evidence from rodent models, including our own, has challenged this view by demonstrating significant neuromuscular impairment in the absence of overt axonal loss^10,12^. These findings have motivated a reexamination of disease mechanisms, particularly focusing on the possibility that primary dysfunction of myelin itself contributes directly to pathogenesis in both CMT1A and HNPP. Given that both disorders result from copy number variation of *PMP22*, a myelin-enriched transmembrane protein with structural similarity to Claudins, and that previous studies have implicated PMP22 in cell adhesion, we hypothesized that PMP22 regulates molecular architecture within peripheral myelin through junctional mechanisms.

Building on prior qualitative reports that localization of E-Cadherin, MAG, and Kv1.1 are altered in *PMP22+/–* and *PMP22– /–* mice, we employed high-resolution confocal imaging and quantitative analysis of teased peripheral nerve fibers from constitutive CMT1A and HNPP mouse models. Our results reveal widespread disruption of adherens junction components at SLIs, including E-Cadherin, β-Catenin, and F-Actin, with divergent structural changes depending on *PMP22* dosage. In CMT1A, these proteins exhibited compacted and fragmented localization, whereas in HNPP, their distribution was broader and mean intensity of F-Actin was elevated. These findings suggest that PMP22 dosage bidirectionally alters junctional organization and SLI morphology, potentially destabilizing Schwann cell cytoskeletal architecture. Interestingly, not all SLI- associated proteins were equally affected. CADM4 remained unchanged, while Connexin29 and MAG showed significant mislocalization in both models, with more dramatic redistribution in CMT1A. Notably, Connexin29 displayed focal accumulations along internodes in CMT1A nerves, suggesting misregulation beyond the SLI compartment. These selective alterations point to protein-specific regulatory effects of PMP22 on myelin architecture and underscore the possibility that PMP22 participates in organizing multiple subdomains of the myelin sheath through distinct molecular mechanisms.

Given the anatomical linkage between SLIs and the Node of Ranvier via the inner mesaxon and known associations between Cx29 and Kv1 channels^19^, we next assessed nodal domain organization. Consistent with a model of impaired axo-glial communication, we observed disrupted patterning of Node of Ranvier axonal components including Nav channels, Caspr, and Kv1.2 channels in both CMT1A and HNPP, with more severe changes frequently observed in CMT1A. These included nodal widening, loss of juxtaparanodal Kv1.2 enrichment, and abnormal spreading of Kv1.2 and Caspr along internodes. Such alterations likely reflect impaired compartmentalization of ion channels critical for saltatory conduction and suggest that junctional instability in myelin has direct consequences for axonal excitability.

Together, our data suggest two pathogenic mechanisms may be working in concert downstream of PMP22 dysregulation: (1) impaired axo-glial communication which likely involves disrupted Schwann cell-mediated metabolic support due to SLI disorganization, and (2) axonal ion channel mislocalization, and consequent axonal ion disequilibrium, due to compromised Node of Ranvier architecture. The former could limit substrate delivery or buffering capacity along long axons, while the latter would degrade conduction efficiency even in the absence of demyelination or axonal loss. These mechanisms are not mutually exclusive and may synergize to impair nerve function. Importantly, our findings also reinforce the notion that PMP22 plays a structural role in maintaining adhesion junction integrity, particularly through regulation of E-Cadherin-based complexes. This is consistent with prior evidence showing a potential direct interaction between PMP22 and E-Cadherin in myelin^16^, and with reports demonstrating that loss of E-Cadherin impairs the formation of autotypic adherens junctions leading to myelin architecture defects^22,23^. The broader disorganization of β- Catenin, F-Actin and potentially p120-Catenin we observed further supports a model in which PMP22 stabilizes adherens junctions that, in turn, maintain subdomain architecture critical for myelin integrity.

This model reframes the pathogenesis of CMT1A and HNPP as diseases rooted in molecular disorganization of myelin architecture, challenging the prevailing view that secondary axonal degeneration is the principal driver of dysfunction. It also provides a mechanistic basis for why compact myelin may appear morphologically preserved while still being functionally impaired. Moreover, the inverse or severity-correlated phenotypes observed in CMT1A and HNPP strengthen the argument that *PMP22* dosage has a direct and bidirectional impact on Schwann cell structure and function. From a therapeutic perspective, these findings highlight junctional complexes, particularly adherens junction components and their cytoskeletal partners, as actionable molecular targets. Strategies aimed at stabilizing or restoring junctional organization could improve conduction properties and metabolic support in CMT1A and HNPP, even in the absence of myelin regeneration. Furthermore, this work offers a conceptual framework for investigating similar junctional pathomechanisms in other inherited dysmyelinating and acquired demyelinating neuropathies.

In summary, we demonstrate that *PMP22* dosage governs the integrity of critical myelin subdomains by regulating adhesion-based molecular architecture. Disruption of this function likely initiates primary myelin instability, leading to impaired metabolic coupling and altered axonal ion homeostasis. These findings advance our understanding of the structural basis of neuropathy in CMT1A and HNPP and open new avenues for therapeutic intervention centered on the stabilization of junctional domains in peripheral nerve.

## Acknowledgements

We thank Regeneron and Lucia Notterpek for sharing the HNPP model mice and Mohamed Farah for providing training with teasing nerve fibers. We also thank Andrew Kowalczyk, Sara Stahley, W. David Arnold, Smita Saxena, Ryan Castoro and Curtis Nutter for advice and helpful scientific discussions and Abigail Chischolm, Hedy Lewis and Sara Tenlep for technical support. We would also like to acknowledge support from the Johns Hopkins Microscope Facility (Office of the Director and the National Institute of General Medical Sciences of the National Institutes of Health award number S10OD023548), the Johns Hopkins Neuroscience Imaging Center and the shared resources available through the University of Missouri Neuro-Muscular Labs and NextGen Precision Health.

## Author Contributions

KRM designed the study, performed experiments and data analysis and wrote the manuscript. MAA and DG performed experiments and data analysis and AH assisted with experimental design and manuscript writing and editing. All authors approved the submitted version of the manuscript.

## Ethics Statement

Animal experiments were conducted with approval from the Johns Hopkins University (approval number Mo21M332) and University of Missouri (approval number 57681) Animal Care and Use Committee. All other experiments were conducted with approval from the University of Missouri Institutional Biosafety Committee (approval number 21140).

## Data Availability Statement

The data that support the findings of this study are available from the corresponding author upon reasonable request.

## References

1. Li, J., Parker, B., Martyn, C., Natarajan, C. & Guo, J. The PMP22 gene and its related diseases. Mol Neurobiol 47, 673–698 (2013).

2. Martyn, C.N. & Hughes, R.A. Epidemiology of peripheral neuropathy. J Neurol Neurosurg Psychiatry 62, 310–318 (1997).

3. Watila, M.M. & Balarabe, S.A. Molecular and clinical features of inherited neuropathies due to PMP22 duplication. J Neurol Sci 355, 18–24 (2015).

4. Attarian, S., Fatehi, F., Rajabally, Y.A. & Pareyson, D. Hereditary neuropathy with liability to pressure palsies. J Neurol 267, 2198–2206 (2020).

5. Park, H.J., Choi, Y.C., Oh, J.W. & Yi, S.W. Prevalence, Mortality, and Cause of Death in Charcot-Marie-Tooth Disease in Korea: A Nationwide, Population-Based Study. Neuroepidemiology 54, 313–319 (2020).

6. Vaeth, S., Vaeth, M., Andersen, H., Christensen, R. & Jensen, U.B. Charcot-Marie-Tooth disease in Denmark: a nationwide register-based study of mortality, prevalence and incidence. BMJ Open 7, e018048 (2017).

7. Vallat, J.M., et al. Ultrastructural PMP22 expression in inherited demyelinating neuropathies. Ann Neurol 39, 813–817 (1996).

8. Krajewski, K.M., et al. Neurological dysfunction and axonal degeneration in Charcot-Marie-Tooth disease type 1A. Brain 123 **( Pt** **7**), 1516–1527 (2000).

9. Manganelli, F., et al. Nerve conduction velocity in CMT1A: what else can we tell? Eur J Neurol 23, 1566–1571 (2016).

10. Moss, K.R., et al. SARM1 knockout does not rescue neuromuscular phenotypes in a Charcot-Marie-Tooth disease Type 1A mouse model. J Peripher Nerv Syst 27, 58–66 (2022).

11. Verhamme, C., et al. Myelin and axon pathology in a long-term study of PMP22-overexpressing mice. J Neuropathol Exp Neurol 70, 386–398 (2011).

12. Robertson, A.M., et al. Comparison of a new pmp22 transgenic mouse line with other mouse models and human patients with CMT1A. J Anat 200, 377–390 (2002).

13. Notterpek, L., et al. Peripheral myelin protein 22 is a constituent of intercellular junctions in epithelia. Proc Natl Acad Sci U S A 98, 14404–14409 (2001).

14. Roux, K.J., Amici, S.A. & Notterpek, L. The temporospatial expression of peripheral myelin protein 22 at the developing blood-nerve and blood-brain barriers. J Comp Neurol 474, 578–588 (2004).

15. Roux, K.J., Amici, S.A., Fletcher, B.S. & Notterpek, L. Modulation of epithelial morphology, monolayer permeability, and cell migration by growth arrest specific 3/peripheral myelin protein 22. Mol Biol Cell 16, 1142–1151 (2005).

16. Guo, J., et al. Abnormal junctions and permeability of myelin in PMP22-deficient nerves. Ann Neurol 75, 255–265 (2014).

17. Hu, B., et al. Tuning PAK Activity to Rescue Abnormal Myelin Permeability in HNPP. PLoS Genet 12, e1006290 (2016).

18. Neuberg, D.H., Sancho, S. & Suter, U. Altered molecular architecture of peripheral nerves in mice lacking the peripheral myelin protein 22 or connexin32. J Neurosci Res 58, 612–623 (1999).

19. Rash, J.E., et al. KV1 channels identified in rodent myelinated axons, linked to Cx29 in innermost myelin: support for electrically active myelin in mammalian saltatory conduction. J Neurophysiol 115, 1836–1859 (2016).

20. Adlkofer, K., et al. Hypermyelination and demyelinating peripheral neuropathy in Pmp22-deficient mice. Nat Genet 11, 274–280 (1995).

21. Fannon, A.M., et al. Novel E-cadherin-mediated adhesion in peripheral nerve: Schwann cell architecture is stabilized by autotypic adherens junctions. J Cell Biol 129, 189–202 (1995).

22. Tricaud, N., Perrin-Tricaud, C., Bruses, J.L. & Rutishauser, U. Adherens junctions in myelinating Schwann cells stabilize Schmidt-Lanterman incisures via recruitment of p120 catenin to E-cadherin. J Neurosci 25, 3259–3269 (2005).

23. Young, P., et al. E-cadherin is required for the correct formation of autotypic adherens junctions of the outer mesaxon but not for the integrity of myelinated fibers of peripheral nerves. Mol Cell Neurosci 21, 341–351 (2002).

24. Schindelin, J., et al. Fiji: an open-source platform for biological-image analysis. Nat Methods 9, 676-682 (2012).

25. Weng, A., et al. Alpha-T-catenin is expressed in peripheral nerves as a constituent of Schwann cell adherens junctions. Biol Open 11(2022).

26. Kourtidis, A., Ngok, S.P. & Anastasiadis, P.Z. p120 catenin: an essential regulator of cadherin stability, adhesion-induced signaling, and cancer progression. Prog Mol Biol Transl Sci 116, 409–432 (2013).

27. Campas, O., Noordstra, I. & Yap, A.S. Adherens junctions as molecular regulators of emergent tissue mechanics. Nat Rev Mol Cell Biol 25, 252–269 (2024).

28. Altevogt, B.M., Kleopa, K.A., Postma, F.R., Scherer, S.S. & Paul, D.L. Connexin29 is uniquely distributed within myelinating glial cells of the central and peripheral nervous systems. J Neurosci 22, 6458–6470 (2002).

29. Meng, X., et al. Necl-4/Cadm4 recruits Par-3 to the Schwann cell adaxonal membrane. Glia 67, 884–895 (2019).

30. Schnaar, R.L. & Lopez, P.H. Myelin-associated glycoprotein and its axonal receptors. J Neurosci Res 87, 3267–3276 (2009).

31. Bao, L. Trafficking regulates the subcellular distribution of voltage-gated sodium channels in primary sensory neurons. Mol Pain 11, 61 (2015).

32. Caldwell, J.H., Schaller, K.L., Lasher, R.S., Peles, E. & Levinson, S.R. Sodium channel Na(v)1.6 is localized at nodes of ranvier, dendrites, and synapses. Proc Natl Acad Sci U S A 97, 5616–5620 (2000).

33. Faivre-Sarrailh, C. Molecular organization and function of vertebrate septate-like junctions. Biochim Biophys Acta Biomembr 1862, 183211 (2020).

34. Marinko, J.T., Kenworthy, A.K. & Sanders, C.R. Peripheral myelin protein 22 preferentially partitions into ordered phase membrane domains. Proc Natl Acad Sci U S A 117, 14168–14177 (2020).

35. Jumper, J., et al. Highly accurate protein structure prediction with AlphaFold. Nature 596, 583–589 (2021).

