## Supplemental Data for "Aberrant Molecular Myelin Architecture in Charcot-Marie-Tooth Disease Type 1A and Hereditary Neuropathy with Liability to Pressure Palsies"

### Supplemental Figures and Legends

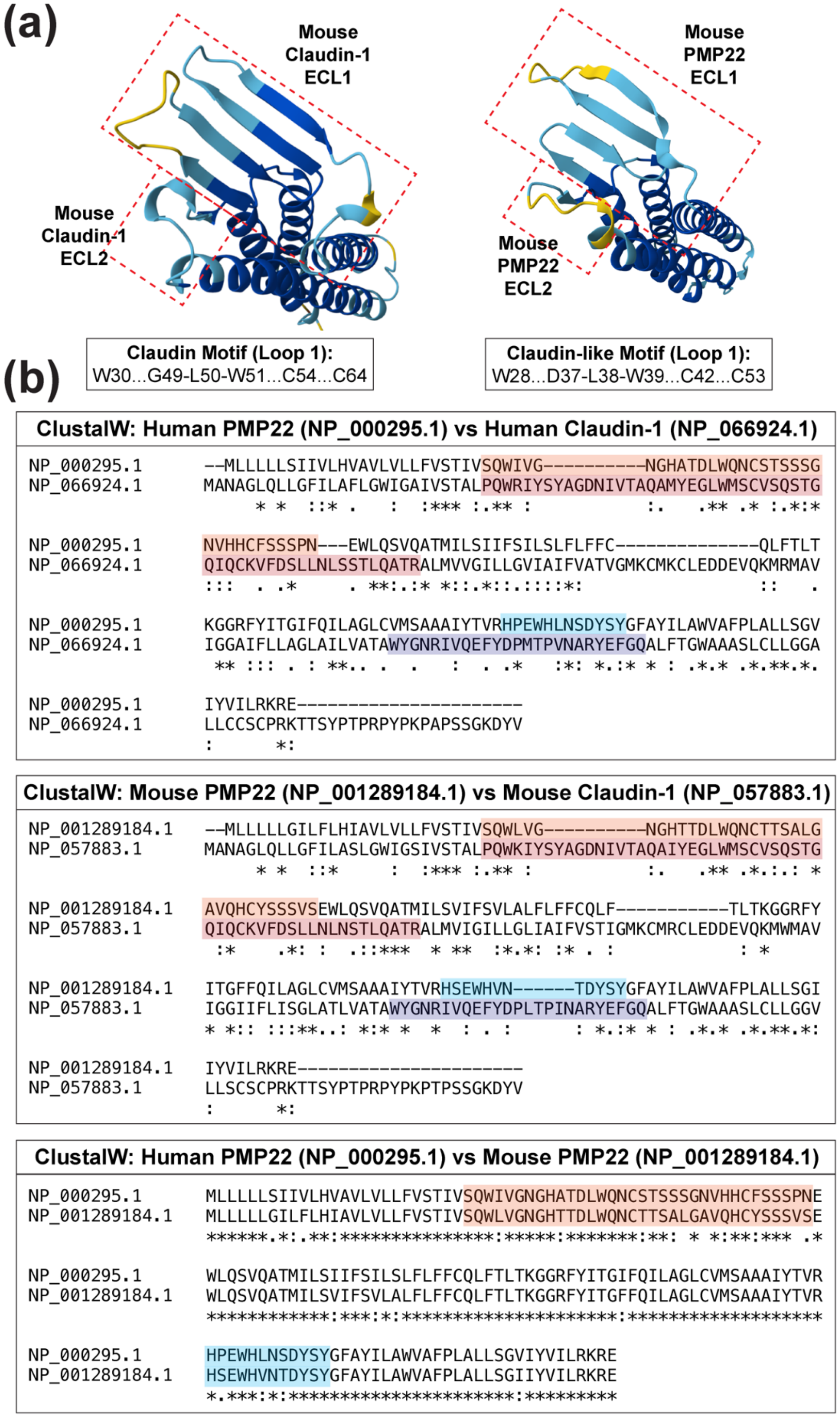


**Supplemental Figure 1. PMP22 Exhibits Structural Similarity to Claudin-1 Despite More Modest Sequence Identity. (a)** AlphaFold predicted structures of mouse Claudin-1 and mouse PMP22 extracellular loops (ECLs) viewed from above (AlphaFold Protein Structure Database). The structural similarity between the

ECLs is striking supporting the notion that PMP22 functions similarly to Claudin proteins. **(b)** ClustalW alignments of human PMP22 with human Claudin-1 (top), mouse PMP22 with mouse Claudin-1 (middle) and human PMP22 with mouse PMP22 (bottom). PMP22 is highly conserved from mouse to human but sequence similarity and identity between PMP22 and Claudin-1 is more limited. The extracellular loop 1 (ECL1) sequences for PMP22 and Claudin-1 are highlighted in orange/red and the extracellular loop 2 (ECL2) sequences for PMP22 and Claudin-1 are highlighted in blue/purple.

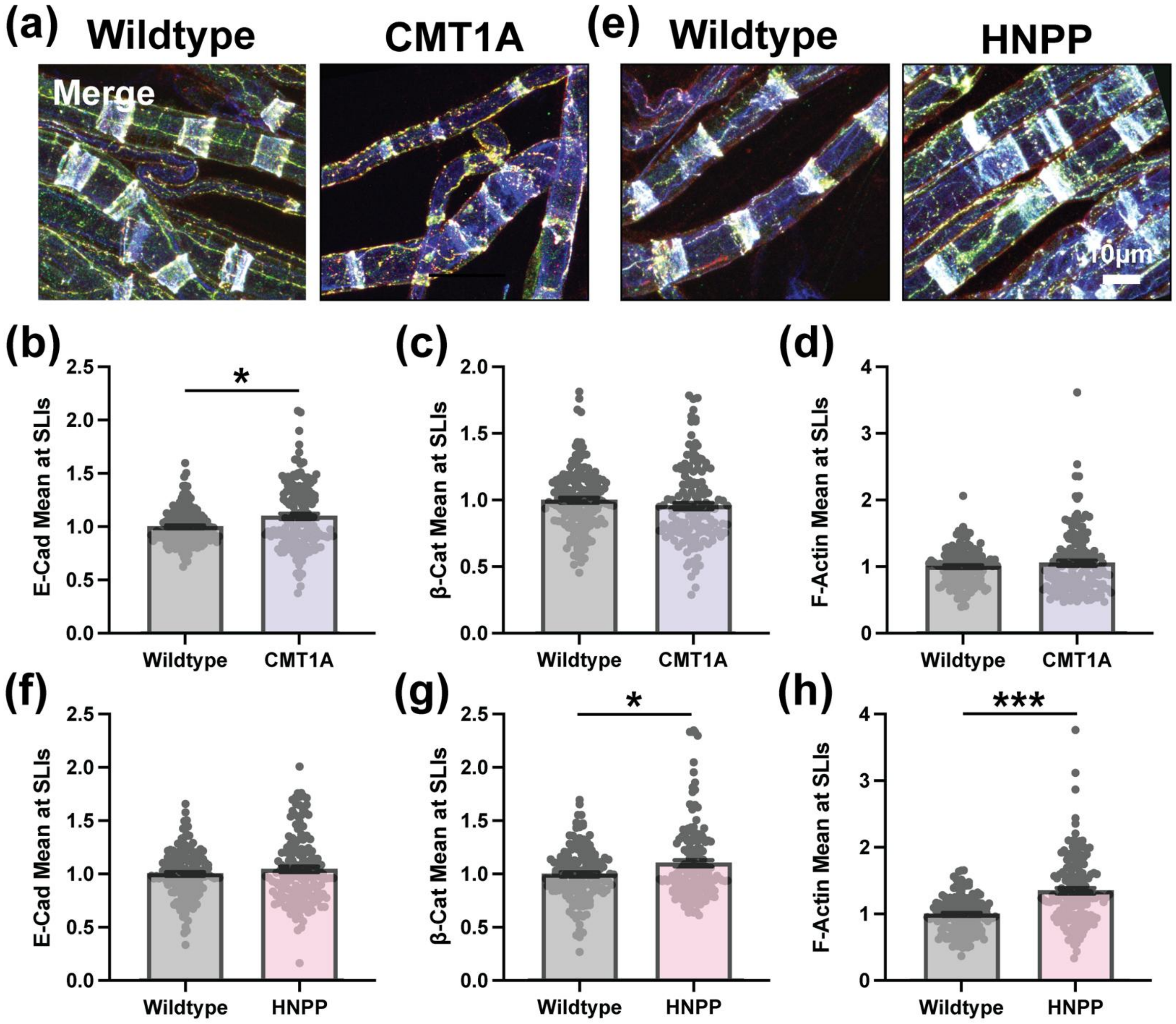


**Supplemental Figure 2. More Subtle Alterations in Adherens Junction and F-Actin Expression at SLIs in CMT1A and HNPP Model Myelin.** Representative images of 3-month-old **(a)** wildtype (WT, C57BL/6J) and CMT1A or **(e)** WT (129S1/SvImJ) and HNPP teased tibial nerve fibers stained for the adherens junction proteins E-Cadherin (red), β-Catenin (green) and F-Actin (blue) merged. Quantification of mean signal intensity at SLIs (readout of protein expression levels/localization) for **(b, f)** E-Cadherin, **(c, g)** β-Catenin and **(d, h)** F-Actin in CMT1A and HNPP, respectively. n=5 animals (~30 SLIs/animal). Bar graphs represent mean ± SEM with individual data points. ***$p<0.05$ demonstrate statistical significance with three separate t-test statistics (unpaired t-test with all individual data points, unpaired t-test with experimental average data points and nested t-test). *$p<0.05$ demonstrate statistical significance with one t-test statistic (unpaired t-test with all individual data points). Scale bar, 10μm.

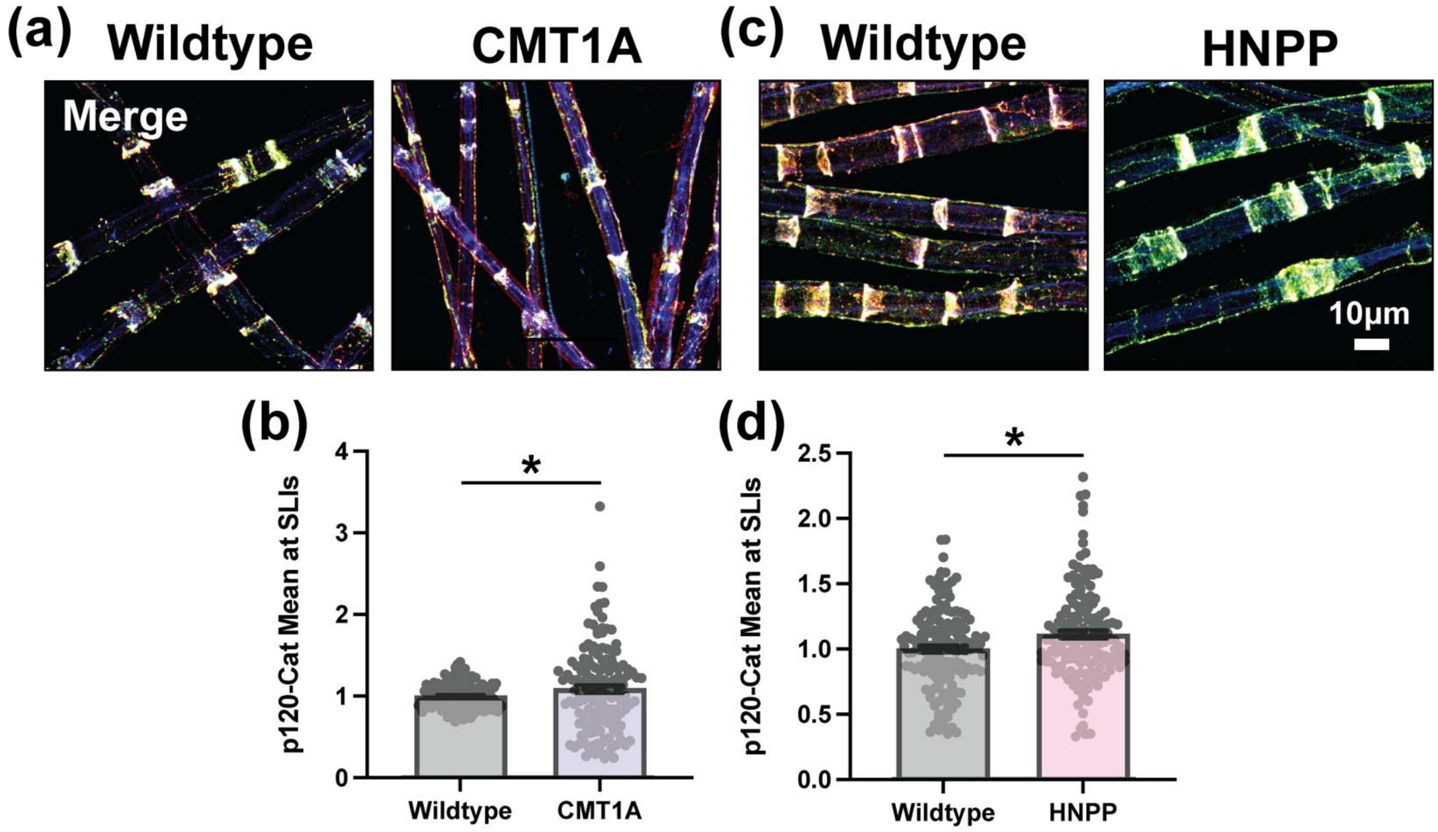


**Supplemental Figure 3. Modest Increase in p120-Catenin Expression at SLIs in CMT1A and HNPP Model Peripheral Nerve Myelin.** Representative images of 3-month-old **(a)** wildtype (WT, C57BL/6J) and CMT1A or **(c)** WT (129S1/SvImJ) and HNPP teased tibial nerve fibers stained for the adherens junction proteins E-Cadherin (red), p120-Catenin (green) and F-Actin (blue) merged. Quantification of mean p120-Catenin signal intensity at SLIs (readout of protein expression levels/localization) in **(b)** CMT1A and **(d)** HNPP. n=5 animals (~30 SLIs/animal). Bar graphs represent mean ± SEM with individual data points. *$p<0.05$ demonstrate statistical significance with one t-test statistic (unpaired t-test with all individual data points). Scale bar, 10μm.

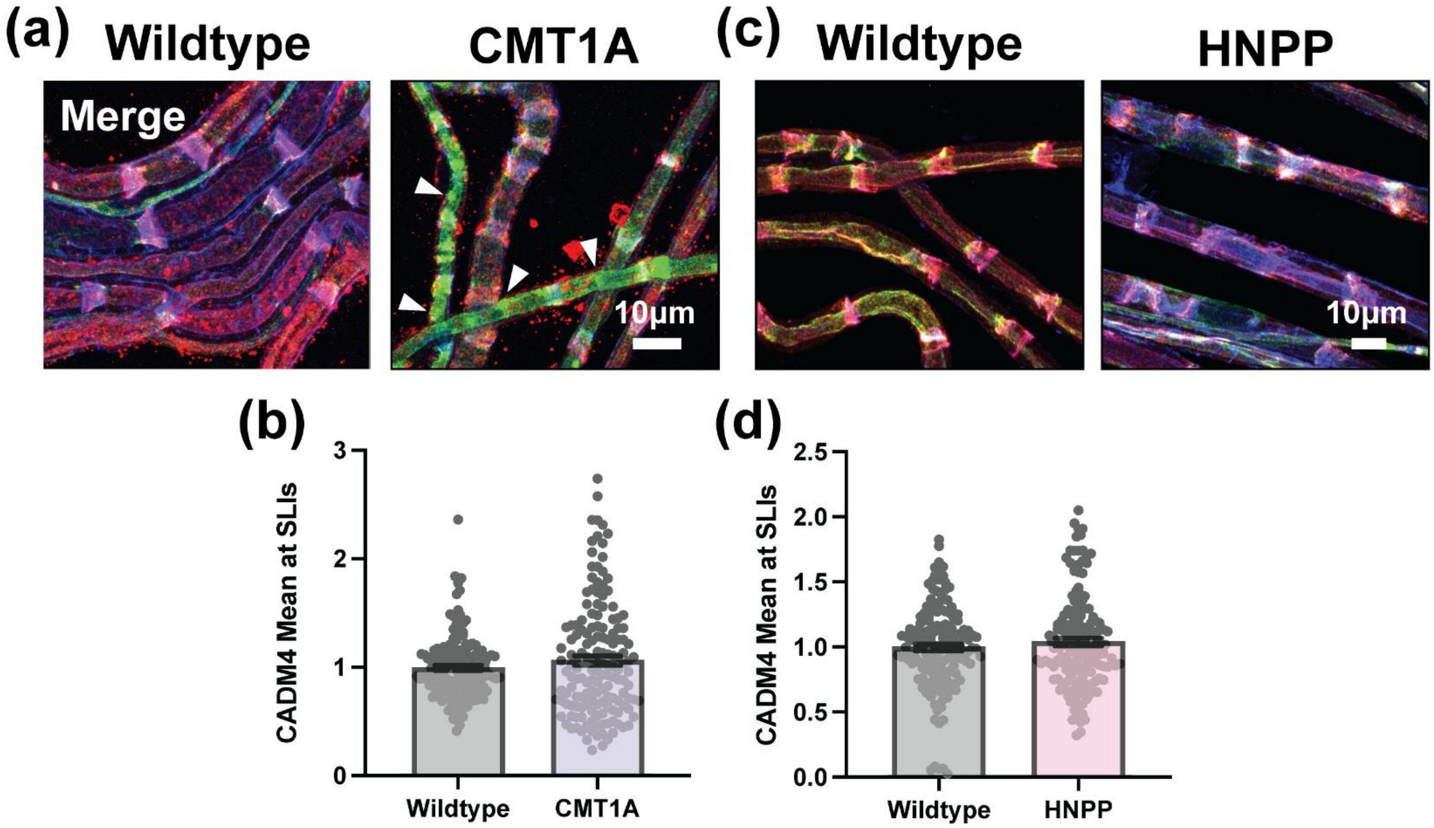


**Supplemental Figure 4. Unchanged CADM4 Expression at SLIs in CMT1A and HNPP Mouse Model Myelin.** Representative images of 3-month-old **(a)** wildtype (WT, C57BL/6J) and CMT1A or **(c)** WT (129S1/SvImJ) and HNPP teased tibial nerve fibers stained for CADM4 (red), Connexin29 (green) and F-Actin (blue) merged. Note the focal accumulations of Cx29 outside the SLI compartment in CMT1A model myelin (arrowheads). Quantification of mean CADM4 signal intensity at SLIs (readout of protein expression levels/localization) in **(b)** CMT1A and **(d)** HNPP. n=5 animals (~30 SLIs/animal). Bar graphs represent mean ± SEM with individual data points. Scale bar, 10μm.

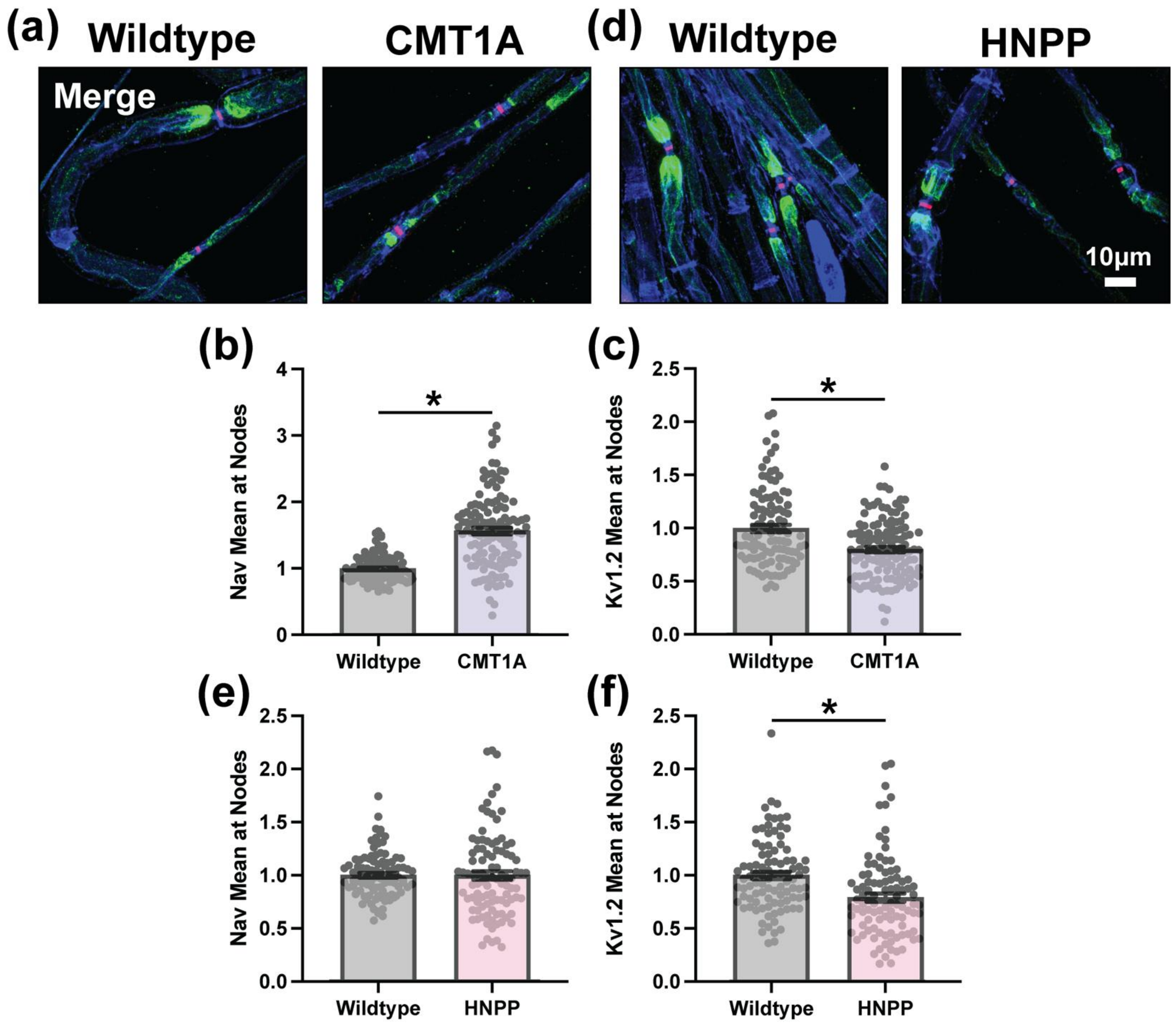


**Supplemental Figure 5. Alterations in Ion Channel Expression at Nodes of Ranvier in CMT1A and HNPP Model Myelin.** Representative images of 3-month-old **(a)** wildtype (WT, C57BL/6J) and CMT1A or **(d)** WT (129S1/SvImJ) and HNPP teased tibial nerve fibers stained with anti-Pan Nav (red), anti-Kv1.2 (green) and Phalloidin to label F-Actin (blue) merged. Quantification of mean signal intensity at Nodes of Ranvier (readout of protein expression levels/localization) for **(b, e)** Nav and **(c, f)** Kv1.2 in CMT1A and HNPP, respectively. n=5 animals (~15-25 nodes/animal). Bar graphs represent mean ± SEM with individual data points. *$p<0.05$ demonstrate statistical significance with one t-test statistic (unpaired t-test with all individual data points). Scale bar, 10μm.

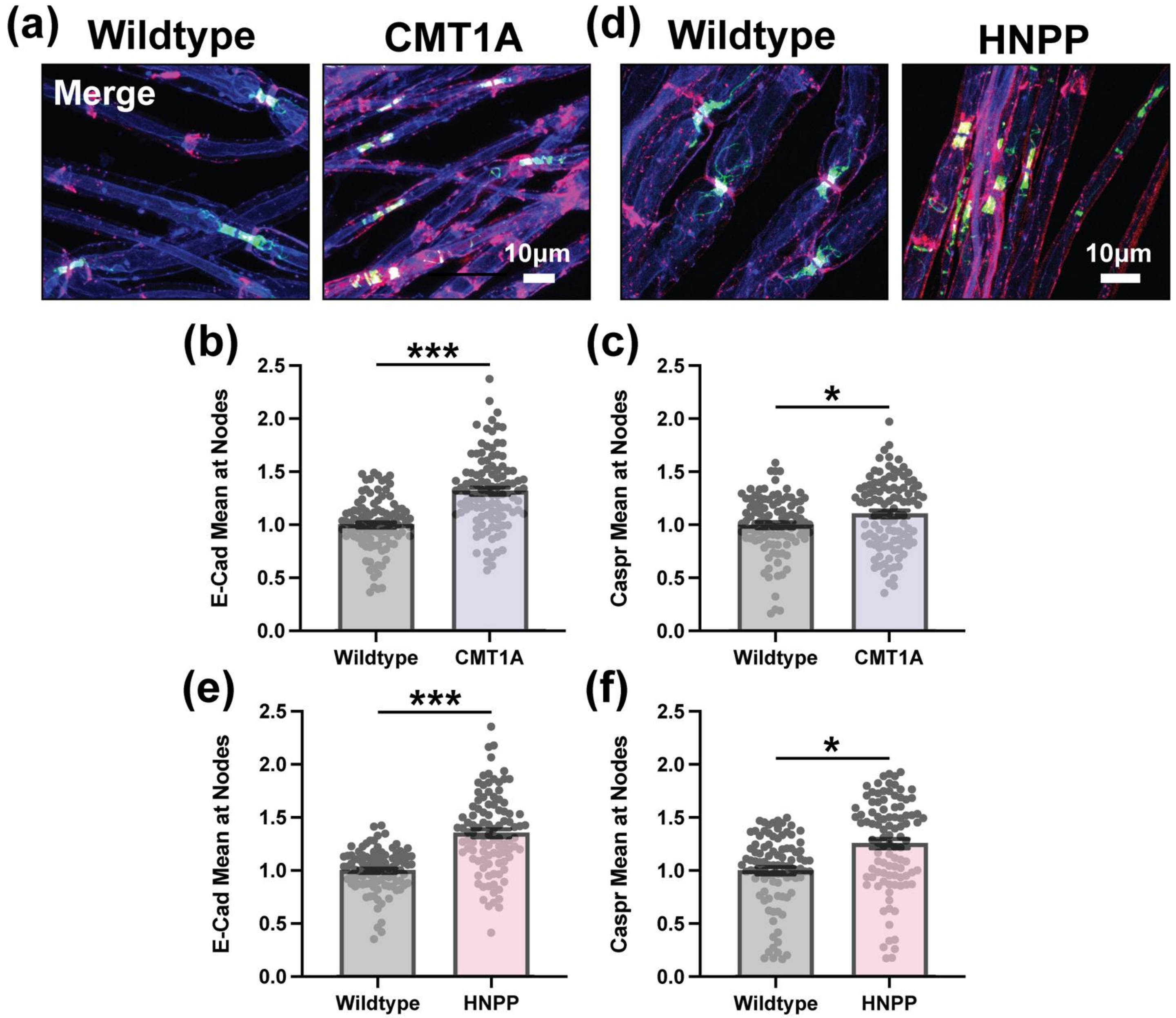


**Supplemental Figure 6. Increased Caspr and E-Cadherin Expression at Nodes of Ranvier in CMT1A and HNPP Model Peripheral Nerve Myelin.** Representative images of 3-month-old **(a)** wildtype (WT, C57BL/6J) and CMT1A or **(d)** WT (129S1/SvImJ) and HNPP teased tibial nerve fibers stained for E-Cadherin (red), Caspr (green) and F-Actin (blue) merged. Quantification of mean signal intensity at Nodes of Ranvier (readout of protein expression levels/localization) for **(b, e)** E-Cadherin and **(c, f)** Caspr in CMT1A and HNPP, respectively. n=5 animals (~15-25 nodes/animal). Bar graphs represent mean ± SEM with individual data points. ***p<0.05 demonstrate statistical significance with three separate t-test statistics (unpaired t-test with all individual data points, unpaired t-test with experimental average data points and nested t-test). *p<0.05

demonstrate statistical significance with one t-test statistic (unpaired t-test with all individual data points). Scale bar, 10μm.
